# Hyperthermia elevates brain temperature and improves behavioural signs in animal models of Autism spectrum disorder

**DOI:** 10.1101/2022.09.20.508692

**Authors:** Ana Belen Lopez-Rodriguez, Carol Murray, John Kealy, Clodagh Towns, Arshed Nazmi, Logan Arnold, Michelle Doran, John Lowry, Colm Cunningham

## Abstract

Autism Spectrum Disorders (ASD) are predominantly developmental in nature and largely genetically determined. There are some human data supporting the idea that fever can improve symptoms in some individuals but the human data for this are limited and there are almost no data to support this from animal models. In the current study we used a whole body hyperthermia (WBH) protocol and systemic inflammation induced by bacterial endotoxin (LPS) to dissociate temperature and inflammatory elements of fever in order to examine the impact of these environmental stressors on behavioural signs in two animal models relevant to ASD: C58BL/6 and Shank3B- mice. While only LPS induced inflammatory signatures in the brain, WBH and LPS induced both overlapping and distinct neuronal cFos activation in several brain regions and modest effects on heat shock gene expression. In behavioural experiments LPS significantly suppressed most activities over 24-48 hours while WBH reduced repetitive behaviours and improved social interaction in C58BL/6 mice. In Shank3B- mice WBH significantly reduced compulsive grooming. The data are the first, to our knowledge, to demonstrate that elevated body temperature, in the absence of underpinning inflammation, can improve some behavioural signs in two distinct animal models of ASD. Given the developmental and genetic nature of ASD, evidence that symptoms may be ameliorated by environmental perturbations indicates that there are possibilities for improving function in these individuals.

## Introduction

Autism is a collection of disorders (Autism Spectrum Disorder, ASD) that is predominantly developmental in nature and largely genetically determined, but there is tentative evidence that environmental changes can impact on symptoms. Fever has long been reported to improve symptoms in individuals with autism. Although this is largely based on carers’ testimony that their children displayed remarkable improvements during febrile episodes, there are some data to support this, with up to 25% of patients displaying evidence of improvement (Curran *et al*., 2007). That study described improvements in repetitive behaviours, stereotypy and irritability and some improvement in speech, that coincided with fever of ≥100.4°F (≥ 38°C). More recent studies have added that carer-reported improvements occur predominantly in those with lower non-verbal cognitive skills and with more repetitive behaviours (Grzadzinski *et al*., 2018). Importantly, given the neuro-developmental nature of ASD, indications that symptoms can improve transiently under certain conditions implies that the circuits affected may possess the structural integrity to perform relatively normally under certain conditions. Understanding the conditions under which these circuits perform optimally offers hope of improving activities of everyday life for those with autism.

Arising from the observation that some individuals with autism function at a higher level than normal during fever, one might speculate that a higher body temperature favours improved circuit function. Increased temperature increases synaptic vesicle release in brain sections (Van Hook, 2020) and spike frequency in the songbird motor pathway (Long & Fee, 2008) and cerebral metabolism has been shown to increase linearly with increased brain temperature (Mrozek *et al*., 2012). In both rats and humans glucose metabolism is altered by hyperthermia in a regionally heterogeneous manner (Nunneley *et al*., 2002).

However during fever, such as that brought about by infection, there is significant production of inflammatory mediators and it is plausible that these mediators, rather than temperature change *per se*, may be the drivers of altered function in the central nervous system, as has been shown in a recent study implicating IL-17a in improving social behaviour in a maternal immune activation (MIA) model of ASD (Reed *et al*., 2020). Equally, it could be that the effects of fever might actually be more impressive if elevated temperature was achieved in the absence of inflammation since there is good evidence that several pyrogenic pro-inflammatory mediators such as IL-1β, IL-6 and prostaglandins also suppress motivation, mood, social activity, arousal and cognition (Saper *et al*., 2012).

Bacterial endotoxin (lipopolysaccharide, LPS) is a pyrogen and can be used to mimic the acute phase of bacterial infection. However LPS experiments must be performed at ambient temperatures that are thermoneutral for mice (i.e. 31°C) in order to produce a 39°C fever in C57BL6 mice (Rudaya *et al*., 2005). Conversely, LPS produces either a mild hypothermia or limited change in body temperature at ambient temperatures of around 22°C (Skelly *et al*., 2013). Therefore, one can use LPS at room temperature to induce systemic inflammation without marked elevation of body temperature. Conversely, whole-body hyperthermia (WBH) can be used to induce fever-range temperatures without triggering the acute inflammation occurring during LPS-induced inflammation (Evans *et al*., 2015). Therefore, we can experimentally dissociate systemic inflammation from fever in order to compare cellular, molecular and behavioural effects of inflammatory and thermal stimuli.

Among the ASD symptoms reported to be alleviated during fever were traits such as hyperactivity, stereotypy, and irritability, all of which could conceivably be suppressed as a result of lethargy and reduced locomotor activity that were likely to be present during the suppressive “sickness behaviour” response of these subjects. Lethargy was described during fever in children with autism (Curran *et al*., 2007) and while suppression of those “hyperactive” behaviours listed would appear beneficial, it is important to establish whether such suppression constitutes a reversal of specific symptoms or simply a suppression of general activity. Therefore, it is important to assess whether fever can boost “negative” features like impaired social approach and anxiety-driven hypoactivity as well as suppressing “positive” features like hyperactivity, repetitive behaviour, excessive grooming, and stereotypy.

In the current study we dissociate fever from its normal inflammatory underpinning and examine its impact on brain and body temperature and on neuronal activation in order to identify what is common among these perturbations and what is distinct. We then use two mouse models relevant to ASD, the C58/J mouse and Shank3B- mice, to assess whether hyperthermia can reverse key deficits in these models.

## Material and Methods

### Animals

Male mice from three different strains were used at 10 weeks old. For initial temperature experiments, C57 mice (C57BL/6J (#000664)) were used for all the treatments and for the assessment of ASD-like models C58 (C58/J (#000669)) and Shank3B- (B6.129-Shank3<TM2GFNG>/J; heterozygous (#017688)) were used. All three strains were from The Jackson Laboratory. Animals were housed in cages of four at 21°C with a 12 h light/dark cycle. Food and water access was *ad libitum*. All animal experimentation was performed under license granted by the Health Products Regulatory Authority, Ireland, with approval from the local ethical committee and in compliance with the Cruelty to Animals Act, 1876 and the European Community Directive, 86/609/EEC. Every effort was made to minimize stress to the animals.

### Subcutaneous temperature transponders implantation

Subcutaneous temperature transponders 14mm wide and 2mm diameter (IPTT-300 (BMDS) were implanted following the manufacturer’s instructions, under light isoflurane anaesthesia, at least three days before being subjected to the WBH protocol. The correct functioning of the transponder was checked before and after the implantation. Temperature monitoring was taken with the IPTT-300 thermoreader (BMDS).

### Brain-implanted thermocouples and real-time temperature recordings

For thermocouple recordings of brain temperature, mice (n=6) underwent stereotaxic surgery to implant MBR-5 intracerebral guide cannulae (ID 457μm, OD 635 μm; BASi Research Products, USA) into the striatum. Mice were anaesthetised with isofluorane (4% for induction, 1.5-3.0 % for maintenance; IsoFlo®, Abbott, U.K.). The surgical site was shaved and disinfected, and lidocaine was administered subcutaneously for local anaesthesia. The skull was exposed, and three stainless steel support screws were implanted into the skull. A burr hole was made over the striatum and the guide cannula was lowered into the caudate putamen (0.3 mm A/P; ±2.0 mm M/L; 3.0 mm D/V), allowing room for the thermocouple to extend into the striatum once inserted into the guide cannula. The guide cannula was cemented into place (Dentalon® Plus, Heraeus-Kulzer, Germany) and, once set, the scalp was sutured to close the wound. All animals were given saline (0.9%; 3 ml/kg body weight) and perioperative analgesia was provided (0.3 mg/kg body weight; Buprecare®, AnimalCare Ltd., U.K.) before animals were allowed to recover in an incubator set to 28 °C, with access to a food gel and hydrogel. Following 7 days of recovery, mice underwent three days of brain temperature recordings in an animal recovery chamber (Vet-Tech. Model: HE010). There were 4-day rest periods between each recording day. On each recording day, the plug was removed from the implanted guide cannula and the thermocouple probe was inserted into the striatum. The thermocouple was an ultrafast T-type implantable device (IT-23; ADInstruments Ltd., UK) which was modified to fit into the implanted MBR-5 guide cannula. Briefly, a 2 cm length of deactivated fused silica tubing (ID 320μm, OD 430μm; Trajan Scientific Europe Ltd., UK) was cut and then carefully inserted under a microscope into a BR microdialysis probe head (BASi Research Products) so that it protruded 7 mm beyond the end of the probe head shaft. A small amount of glue (WEICON Epoxy Minute Adhesive) was then applied to the end of the shaft to fix the silica in place. After ca. 1 h the thermocouple was inserted through a 1.8 cm length of PEEK tubing (ID 860μm, OD 1270μm; Plastics One, Roanoke, VA, USA) which was pushed well up the probe so that it was out of the way until later gluing. The thermocouple was then carefully inserted into the silica tubing under a microscope until it protruded ca. 1 cm. A small amount of epoxy glue was applied to the end of the silica and the thermocouple gently pulled back until it was 1 mm from the end of the silica as confirmed using a digital calliper. This was then left to dry for ca. 1 h before placing some epoxy glue around the silica at the top of the BR probe head. The PEEK tubing was then immediately pulled down and carefully inserted into the epoxy. Following overnight storage, the modified thermocouple was connected to a T-Type Thermocouple Pod (ADInstruments Ltd.), which was in turn connected to the pod port of an e-Corder (eDAQ Pty Ltd, Australia), and the operational characteristics (temperature vs. voltage output) tested using the suppliers’ guidelines (ML312 T-type Pod Manual, ADInstruments Ltd.). This involved placing the thermocouple in a jacketed cell (ALS Ltd, IJ Cambria Scientific Ltd, Llanelli, UK) attached to a thermostatically controlled circulating water bath (Julabo Corio CD-BC4, Fisher Scientific, Dublin, Ireland), and recording the temperature vs. voltage output in Chart TM (Version 5, eDAQ Pty Ltd) at a sampling rate of 1 Hz. Raw temperature recordings in vivo were made in Chart TM (Version 5, eDAQ Pty Ltd, Australia) at a sampling rate of 10 Hz. On each recording day, baseline recordings of brain temperature were made for 1 hour. Following the baseline recordings, the mice went through one of three different interventions: (a) Day 1: Room temperature protocol. (b) Day 5: Whole body hyperthermia (WBH) protocol. (c) Day 9: LPS protocol (250mg/Kg; i.p.). During brain temperature recording experiments, core body temperatures were measured using the subcutaneous temperature transponders (IPTT-300 (BMDS) and the IPTT-300 thermoreader (BMDS) every 20 minutes without removing the mouse from the test chamber. All activity was marked on the real-time brain temperature recording.

### Whole Body Hyperthermia and LPS treatment

Mice were exposed to whole body hyperthermia (WBH) for 4 hours by transferring them to a small animal recovery chamber (Vet-Tech. Model: HE010) set at 38.5°C (30±1% humidity) to reach the target body temperature of 39.5 ± 0.5 °C. This was adapted for C58 strain, see results section. WBH protocol was invariably performed at the same time of day (8 a.m.) to minimize the effects of the circadian rhythm that occurs in the body temperature of rodents. Prior to starting the WBH protocol, the animals were transferred from their home cage to an empty new cage without bedding, food, or water to allow them to habituate to a new space (T_-1h_) and their body temperature and weight were taken. One hour later (T_0h_), their body temperature was recorded again using the thermoreader and subcutaneous temperature transponder (IPTT-300, BMDS) and they were injected with either saline for controls and WBH groups or LPS (250µg/kg) before being transferred to the WBH chamber or the room temperature (RT) chamber (21±1°C; 50±1% humidity). They were left to freely move and explore, and their temperatures were taken every 20 minutes from T_0h_ to T_4h_. Two hours after starting WBH (T_2h_) the temperature and weight of the animals was taken and every animal, independent of treatment, was injected with 500µl of saline to avoid dehydration and quickly returned to the chamber. Temperature recording every 20 minutes was resumed. At 4 hours, temperature and weights were taken and every animal, independent of treatment, was again injected with 500µl of saline. Although some WBH animals did have short episodes of jumping in the heating chamber, they were largely lethargic for the 4 hours period and, on removal from the heating chamber, did rapidly re-establish normal locomotor behaviour, while LPS animals remained lethargic for many hours. For molecular experiments, animals were euthanized immediately after the WBH protocol and for behavioural studies, they underwent a ‘step-down’ cooling protocol to avoid the rebound hypothermia that occurs after WBH (Wilkinson *et al*., 1988). The ‘step down’ protocol was as follows: the heated animals were returned to the WBH chamber, which was set, sequentially, at 32, 28, 23°C and RT, for 20 minutes at each point in the sequence. Their temperature was also monitored and after the step-down protocol they were either returned to their home cage or started a panel of behavioural tasks (T_5h_), depending on the requirements of the experiment. (See figure 1 for protocol schematic).

**Figure 1.**
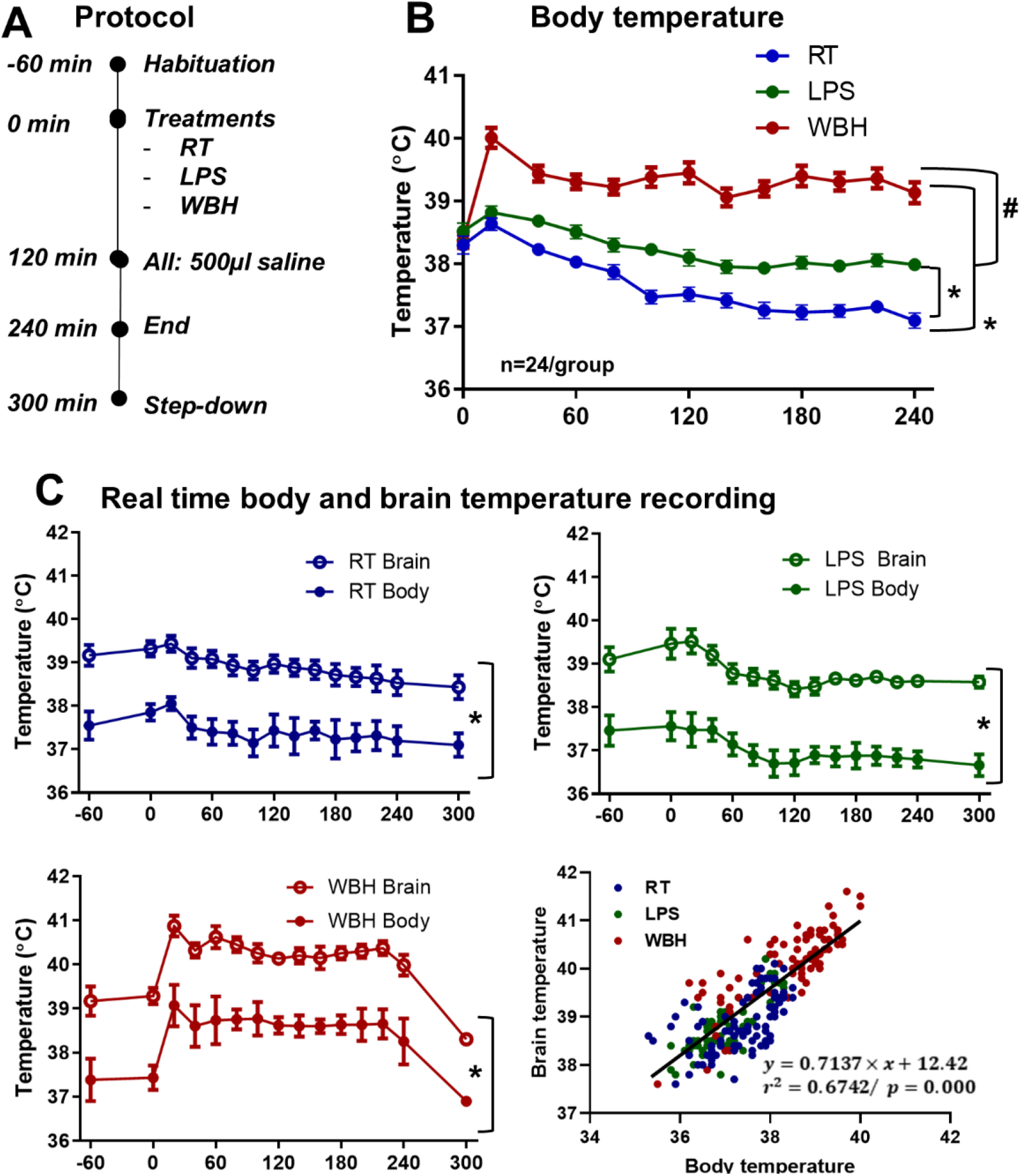
Whole-Body Hyperthermia produces elevation of body and brain temperature. A) Schematic timeline of the whole-body hyperthermia (WBH) protocol. B) Body temperature time course (°C) over the five hours duration of the protocol, measured by subcutaneous temperature transponders. Data are shown as Mean ± SEM (n=24 for each group) and are analysed by repeated measures two-way ANOVA (* vs. RT group; # vs. LPScvf. Bonferroni post-hoc, p<0.05 °C). C) Real time body and brain temperature (°C) over the five hours duration of the protocol. Brain temperature was monitored using brain-implanted thermocouples in a small subset of those animals monitored by sub-cutaneous transponders (n=5 for LPS and 6 for other groups). Abbreviations: RT (room temperature); LPS (intraperitoneally injected lipopolysaccharide, 250 µg/kg); WBH (whole-body hyperthermia).

### Tissue preparation

For the analyses of transcriptional changes, animals were terminally anesthetized with sodium pentobarbital at 4 h post WBH protocol (Euthatal; Merial Animal Health) and rapidly transcardially perfused with heparinised saline before the dissection of hypothalamus, hippocampus and amygdala that were snap frozen in liquid nitrogen and stored at −80°C until use. Animals for immunohistochemical examination were terminally anesthetized with sodium pentobarbital (Euthatal; Merial Animal Health) and transcardially perfused with heparin-saline followed by 4% paraformaldehyde (PFA). Brains were gently removed and postfixed in 4% PFA. Coronal sections, 50 μm thick, were obtained using a Vibratome (Leica, Laboratory Instruments and Supplies, Ashbourne) to perform immunohistochemistry.

### RNA extraction, cDNA synthesis and quantitative PCR

Total RNA was isolated using the RNeasy Plus Mini method (Qiagen, Limburg, Netherlands) following the manufacturer’s instructions. The RNA yield and quality of each sample were quantified based on Optical Density (OD) using the NanoDrop’ND-1000 UV–vis spectrophotometer (Thermo Fisher Scientific). cDNA synthesis was carried out using a High Capacity cDNA Reverse Transcriptase Kit (Applied Biosystems, Warrington, UK). Primer and probe sets were designed using NCBI Nucleotide tool and amplified a single sequence of the correct amplicon size, as verified by SDS-PAGE. Primer pair/probe sequences are shown in Table 1. Samples for RT-PCR were run in duplicate using FAM-labelled probes or SYBR green dsDNA-intercalating fluorescent dye (Roche) in a StepOne Real-Time PCR system (Applied Biosystems, Warrington, UK) under the cycling conditions: 95°C for 10 min followed by 95°C for 10 secs and 60°C for 30 secs for 40-45 cycles. Quantification was achieved by exploiting the relative quantitation method. We used cDNA, prepared from isolated RNA that was pooled from the brains of WBH-treated and LPS-injected mice as a standard that expressed all genes of interest. Serial 1 in 4 dilutions of this cDNA were prepared in order to construct a linear standard curve relating cycle threshold (CT) values to relative concentrations, as previously described (Cunningham *et al*., 2005). Gene expression data were normalized to the housekeeping gene 18S and expressed as relative concentration.

**Table 1.**
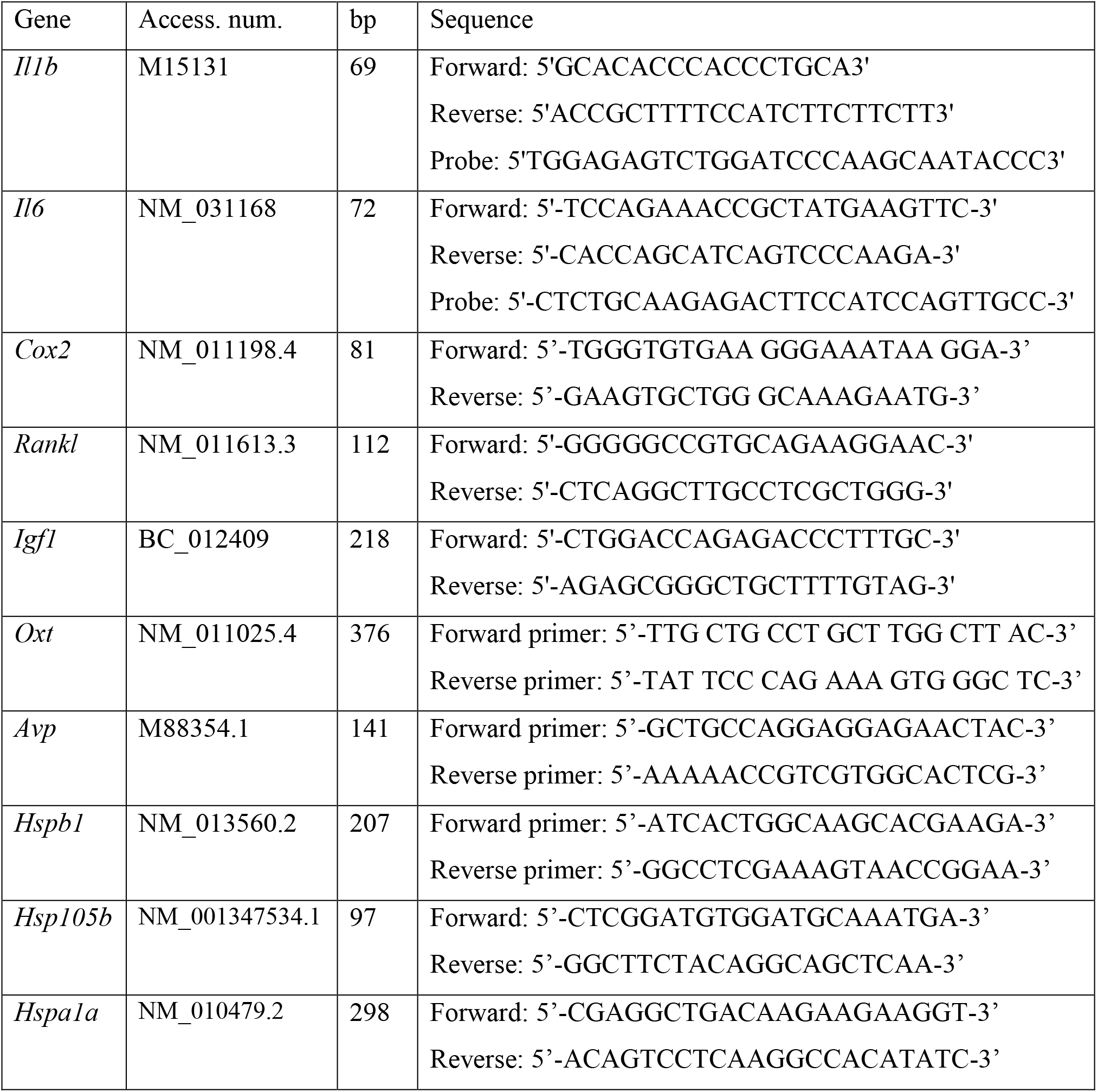
Primer and probe sequences. Primer and probe sequences for quantitative PCR. All probes used FAM as reporter. Where probe sequence is not shown these assays were performed using SYBR green.

### Immunohistochemistry

Immunohistochemistry was carried out on free-floating sections under moderate shaking. All washes and incubations were done in 0.1M phosphate buffer pH 7.4, containing 0.3% bovine serum albumin and 0.3% Triton X-100. The endogenous peroxidase activity was quenched in a solution of 3% hydrogen peroxide in 30% methanol. Sections were incubated overnight at 4 °C with anti-c-Fos (C-10) mouse monoclonal IgG2, (Santa Cruz Biotechnology, Heidelberg, Germany), diluted 1:1000 in the presence of 5% normal horse serum (Vector Laboratories inc, Burlingame, CA), Next day, sections were incubated for 2 hours with biotinylated horse anti-mouse secondary antibody (1:300, Vector). After several washes in phosphate buffer saline (PBS), ABC method was used (Vectastain Kit, PK6100 Vector) and the reaction product was revealed using 3, 3’ diaminobenzidine as chromogen (Sigma Aldrich) and H_2_O_2_ as substrate. Finally, sections were dehydrated, mounted on gelatinized slides, coverslipped, and examined and photographed using an Olympus DP25 camera (Mason) mounted on a Leica DM3000 microscope (Laboratory Instruments and Supplies, Ashbourne), captured using CellA™ software (Olympus, Mason).

### cFOS analyses

The assessment of cFos activation was by a qualitative analysis using series of sections separated by 300 µm, beginning from the olfactory bulbs and continuing to the brain stem. Microscope images were taken at 10x and 20x and scored by the investigator blinded to treatment. Every section in which a given region of interest was present was scored (1 to 4) and averaged within the same animal, and then averaged within the treatment group (n=5-6). The score corresponds to: + minimal activation, ++ medium activation, +++ high activation. The regions of interest used to capture the two main components of fever: temperature and inflammation were chosen based on previous works that analysed cFos activation under cold/warm temperature protocols (Bratincsák & Palkovits, 2004) and after LPS treatment (Lacroix & Rivest, 1997).

### Blood glucose measurements

Two different methods were used to assess blood glucose. For serial sampling, the blood glucose levels were measured via serial tail vein microsampling (less than 10μl) one hour before starting the protocol and then at 40 mins, 2h, 4h, 7h and 24h. Animals were bled using a 30G lancet and glucose was determined in the blood drop with the precision Xtra glucometer (Abbott). In a different set of experiments, the glucose levels were determined in the first drop of blood from the right atrium immediately before performing transcardial perfusion (4h or 24h after the heating protocol).

### Plasma ELISA assays

Animals were terminally anaesthetised at 4 hours post WBH protocol with sodium pentobarbital (Euthatal, Merial Animal Health). The thoracic cavity was opened, and blood collected in heparinised tubes directly from the right atrium of the heart. This whole blood was spun at 1.5 x g for 15 minutes to remove cells; the plasma was then aliquoted and stored at −20°C until use. These samples were then diluted appropriately and analysed for IL-1β, TNF-α and IL-6 by sandwich-type ELISA, using ELISA MAX Mouse IL-6, ELISA MAX Mouse IL-1β (Biolegend, San Diego, US), and DuoSet Mouse TNF-α (R&D Systems, Minneapolis, US). The required capture and detection antibodies, cytokine standard and Avidin-HRP (IL-6) or Streptavidin-HRP (TNF-α) were supplied with each respective kit, however for IL-1β a more sensitive Streptavidin poly-HRP (Sanquin, Amsterdam, The Netherlands) was used in place of the supplied one. Optical density was read at 450 nm with correction at 570 nm. Standard curves for each antibody were used and samples were quantified only if the absorbance fell on the linear portion of the standard curve. Reliable quantification limits for the assays used were IL-1β 31.25 pg/ml, TNF-α 15.6 pg/ml and IL-6 15.6 pg/ml.

### Behavioural tasks

#### Burrowing

Burrowing is a species typical behaviour which may be analogous to domestic behaviours and the task was performed following Deacon’s protocol (Deacon, 2006, 2009). Burrows were made from a 200mm long, 68mm diameter black plastic tube. One end of the tube was closed with a piece of plastic from the same material and the other end was open and raised 30mm above the cage floor by two 50mm screws. Animals were placed into individual opaque cages with fresh bedding and provided with water and the burrowing tube filled with 300g of food pellets as substrate. The food pellets remaining in the burrowing tubes were measured after 2 hours and at 24 hours and this weight was subtracted from 300g to calculate the burrowing activity for each mouse. At 24 hours, the mice were returned to their home cages.

#### Open field activity

The open field test was used to assess spontaneous activity in a novel environment, and it also served as the habituation period for the social interaction test that immediately followed it (detailed below). Briefly, mice were allowed to freely and individually explore an open field arena (58 × 33 × 19 cm; divided into squares of 10 × 10 cm) for 10 min. Activity was monitored via an overhead camera and recorded using AnyMaze software (version 4.99). The mice were assessed on parameters such as distance travelled, mean speed, rotations, time spent in the outer and inner zone and time freezing.

#### Social interaction

We assessed social interaction using a rectangular arena 58 × 33 × 19 cm. Two wire mesh cages (9 cm diameter) were placed in the middle of each half of the arena. The test began with a habituation period to the arena for 10min, (this was used to take measurements of locomotor activity as described in ‘open field activity’ above). Thereafter the mouse was allowed to adapt, for 5 minutes to the placement of two small empty wire mesh cages for 5 min. Finally, the social preference test was conducted for 10 min, during which an unfamiliar (stranger) mouse of the same age, weight and sex was placed into one of the mesh cages whereas the other mesh cage contained an inanimate object. AnyMaze software was linked via an overhead camera and used to recorded test mice, tracking the movement of the test mice, and recording parameters such as time spent sniffing the novel mouse and the number of sniffs, time spent in the area of the mouse mesh cage vs. the empty cage and time freezing. After the trial, the arena and wire cages were cleaned with 10% ethanol.

#### Backflips and Upright Scrabbles

Backflips and upright scrabbles have been described as specific behavioural alterations of the C58/J strain (Moy *et al*., 2008, 2014; Muehlmann *et al*., 2012; Blick *et al*., 2015; Whitehouse *et al*., 2017). The test has two parts. First, a period (5 min) of adaptation to an opaque plastic cage (19.5 x 31 x 13 cm) with a regular metal bar lid placed on top provided with fresh bedding for each mouse. The second part consisted of observation and counting of the number of backflips and upright scrabbles for a period of 30 min. Back flipping manifests as backward somersaulting, often with the assistance of the cage lid, while upright scrabbling consisted of rapidly running or climbing ‘on the spot’, usually against a wall or corner, which may be related to wall-climbing stereotypy.

#### Marble burying

An opaque plastic cage of 45 (l) × 23 (w) × 13 (h) cm was used for this test. The cage was filled approximately 6-8cm deep with wood chip bedding, lightly patted down to flatten the surface and make it even. A regular pattern of 20 glass marbles was placed on the surface: 5 columns 8 cm apart and 4 rows 4 cm apart. The mouse was placed in the cage and left for 30 min. After that time, the marbles that were completely buried or buried to 2/3 their depths were counted.

#### Grooming

One of the specific behavioural alterations of the Shank3B- strain is excessive, sometimes injurious grooming (Peça *et al*., 2011; Mei *et al*., 2016; Balaan *et al*., 2019). To assess this, mice were placed individually in a clean clear plastic cage with fresh bedding and the lid, and they were left for 5 min to adapt to the new environment. After that, the time spent grooming was measured over 5 min.

#### Elevated Zero Maze

Shank3B-mice have been shown to spend less time in the open arms during the Elevated-Zero-Maze (EZM) (Peça *et al*., 2011; Mei *et al*., 2016). The EZM is a modification of the plus-maze based on two conflicting innate tendencies: exploring a novel environment and avoiding elevated and open spaces that constitute a risky situation. The EZM consists of an elevated circular track divided into 4 equal lengths: 2 lengths of track enclosed by an opaque wall on both sides and 2 equal lengths that do not have a surrounding wall. Open and closed areas alternate. Mice were placed individually into one of the closed arms and left to explore for 5 min. The time spent in exploring enclosed versus open arms, the latency to enter the open arms and the number of risk-assessment events were counted.

#### Horizontal Bar

Shank3B- mice present mild motor abnormalities that are increased in Shank3KO (Peça *et al*., 2011; Wang *et al*., 2011; Mei *et al*., 2016). The horizontal bar test was used to assess the muscular strength, motor coordination and prehensile reflex. This test consisted of a 26 cm long metal bar, 0.2 cm diameter, supported by a 19.5 cm high column at each end. Each mouse was held by the tail, placed with its front paws at the central point of the bar, and rapidly released. A score was assigned depending on whether and when the mouse fell, whether it held on for 60 seconds, or whether it reached a supporting column. Animals score 1 if they fell off within 10 seconds, score 2 if they held on for 11-59 seconds, score 3 if they held on for 60 seconds or reached the safe platform in 60 seconds, score 4 if they reached the safe platform within 30 seconds and score 5 if they reached the platform within 10 seconds.

#### Area under the curve

Several parameters were compared in WBH versus RT groups using an area under the curve (AUC) calculation. The trapezoidal rule method (Curry & Whelpton, 2016) was used to calculate the AUC (*A*): 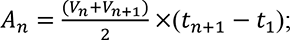 where *V* is the study variable and t is the time (from 0h to 48h). The obtained values were then plotted in two columns and analysed by an unpaired two-tailed t-test or the Mann-Whitney test if data did not pass the assumptions for parametric analyses.

#### Statistical analyses

As indicated in table 2, multiple groups were analysed by repeated measures, two-way, analysis of variance (ANOVA), with factors being treatment (room temperature (RT), LPS or whole-body hyperthermia (WBH)) and time (0h to 48h). Post hoc comparisons (Bonferroni’s test) were performed with a level of significance set at p <0.05. For the analyses of non-repeated measures, one-way analysis of variance (ANOVA) was performed to compare RT, LPS and WBH. Data were not always normally distributed, and, in these cases, nonparametric tests were used (Kruskal–Wallis and post hoc pair-wise comparisons with Mann–Whitney U-test). Post hoc comparisons were performed with a level of significance set at p <0.05. For two-group comparisons, data were analysed using unpaired two-tailed t-test when they were normally distributed, and the Mann-Whitney test was run if data did not pass the assumptions for parametric analyses. For correlation analyses, data from all groups were pooled and adjusted by a linear regression model. Data are presented as mean ± standard error of the mean (SEM). Symbols in the graphs denote post-hoc tests. Statistical analyses were carried out with the SPSS 22.0 software package (SPSS, Inc., Chicago, IL, USA).

**Table 2.**
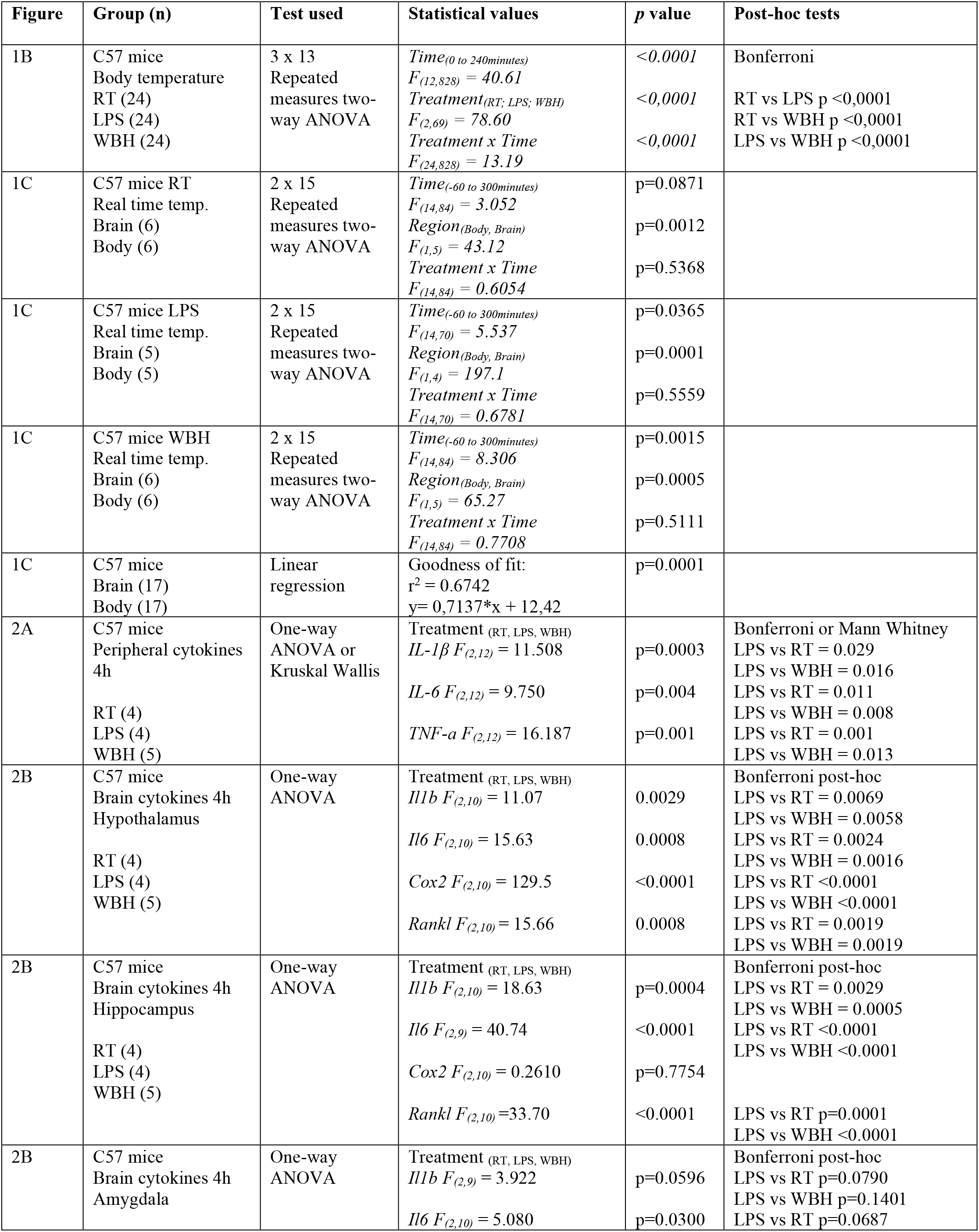
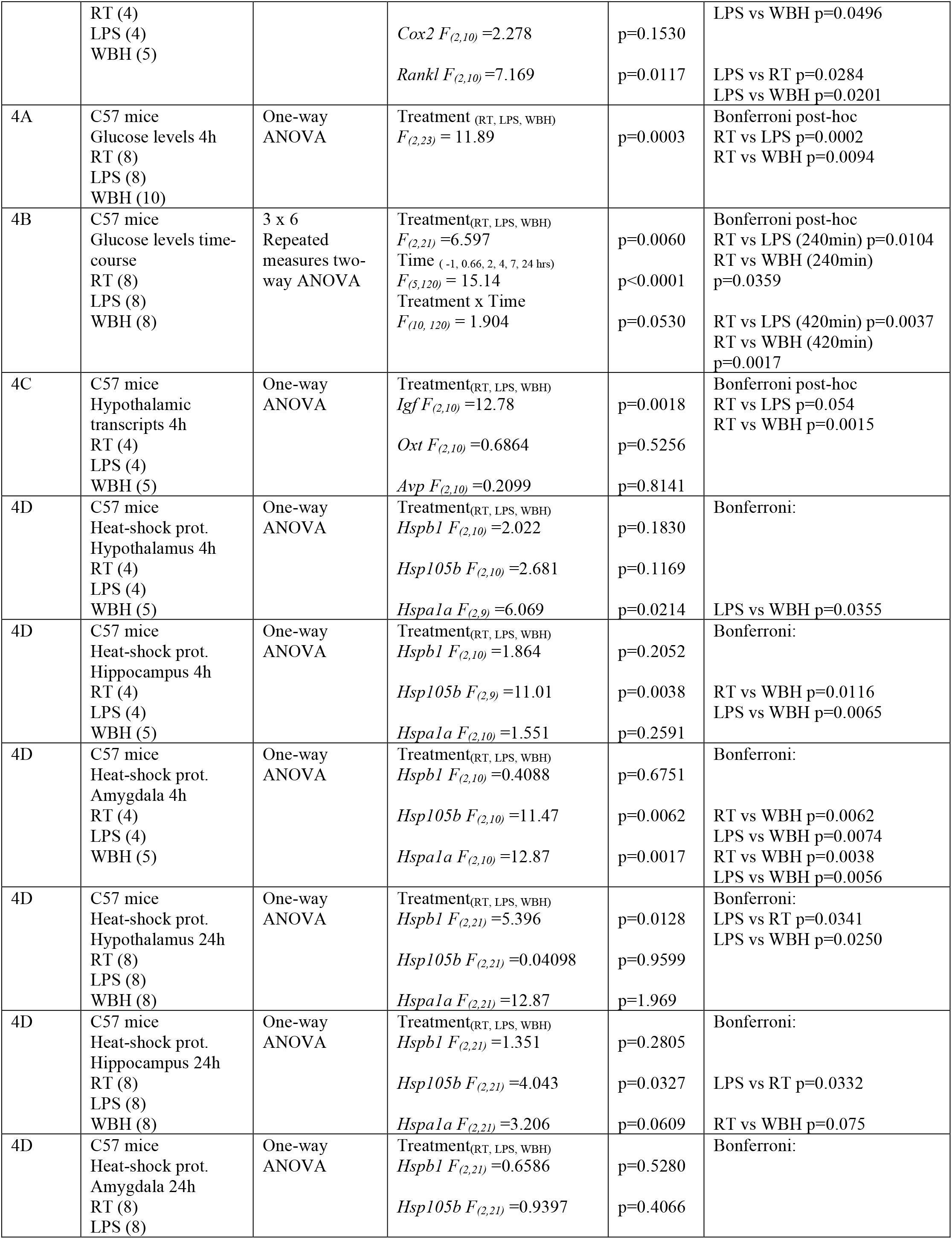
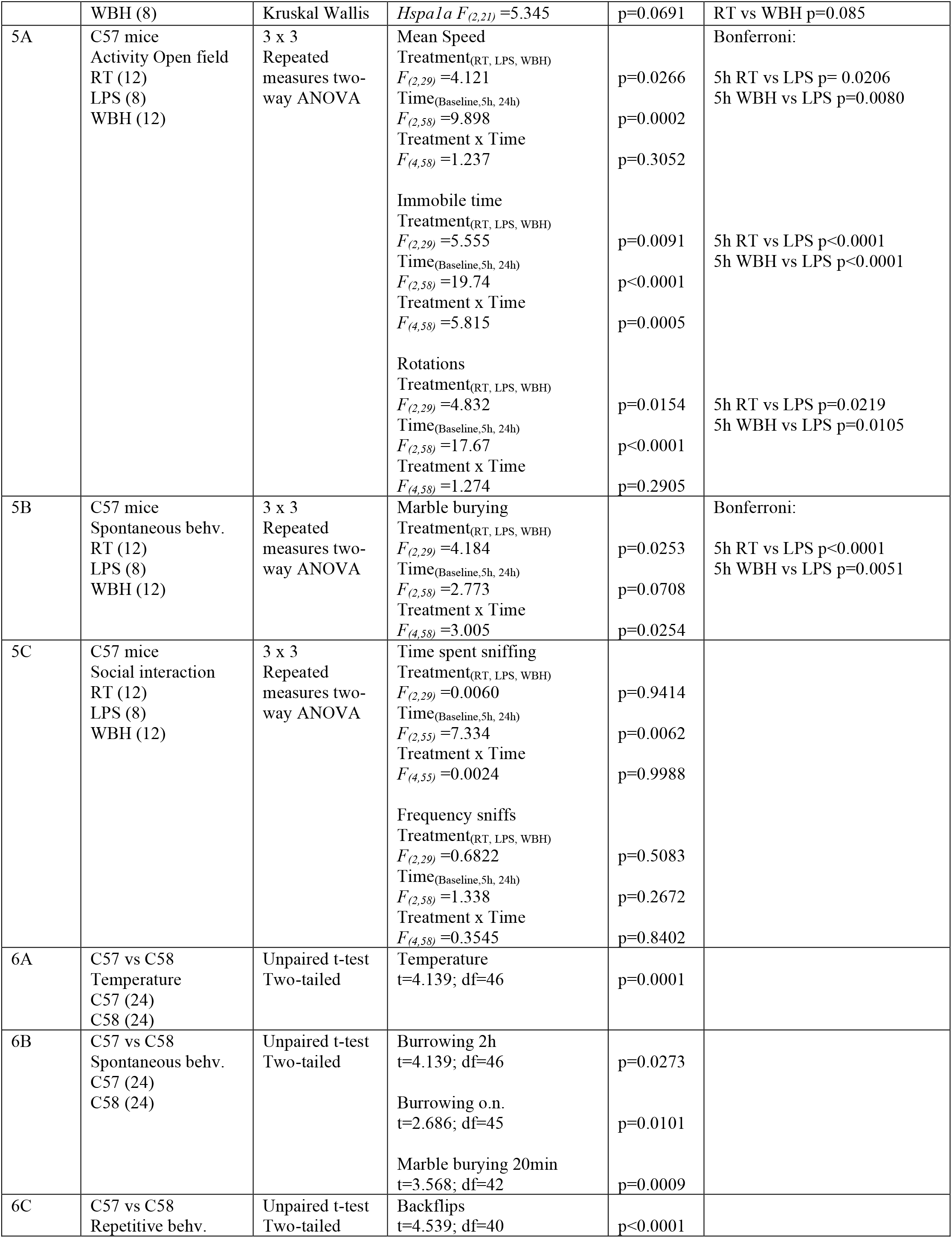
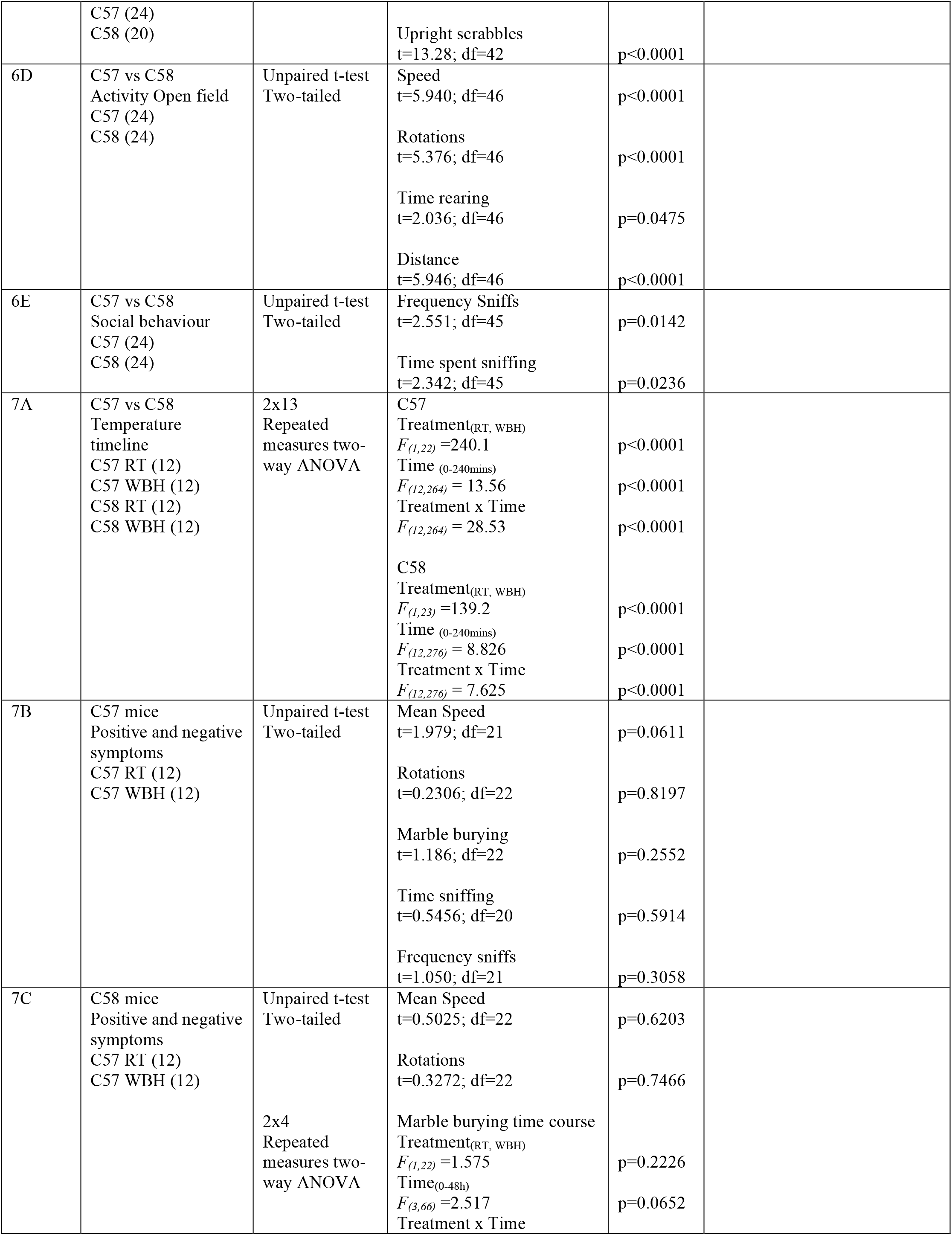
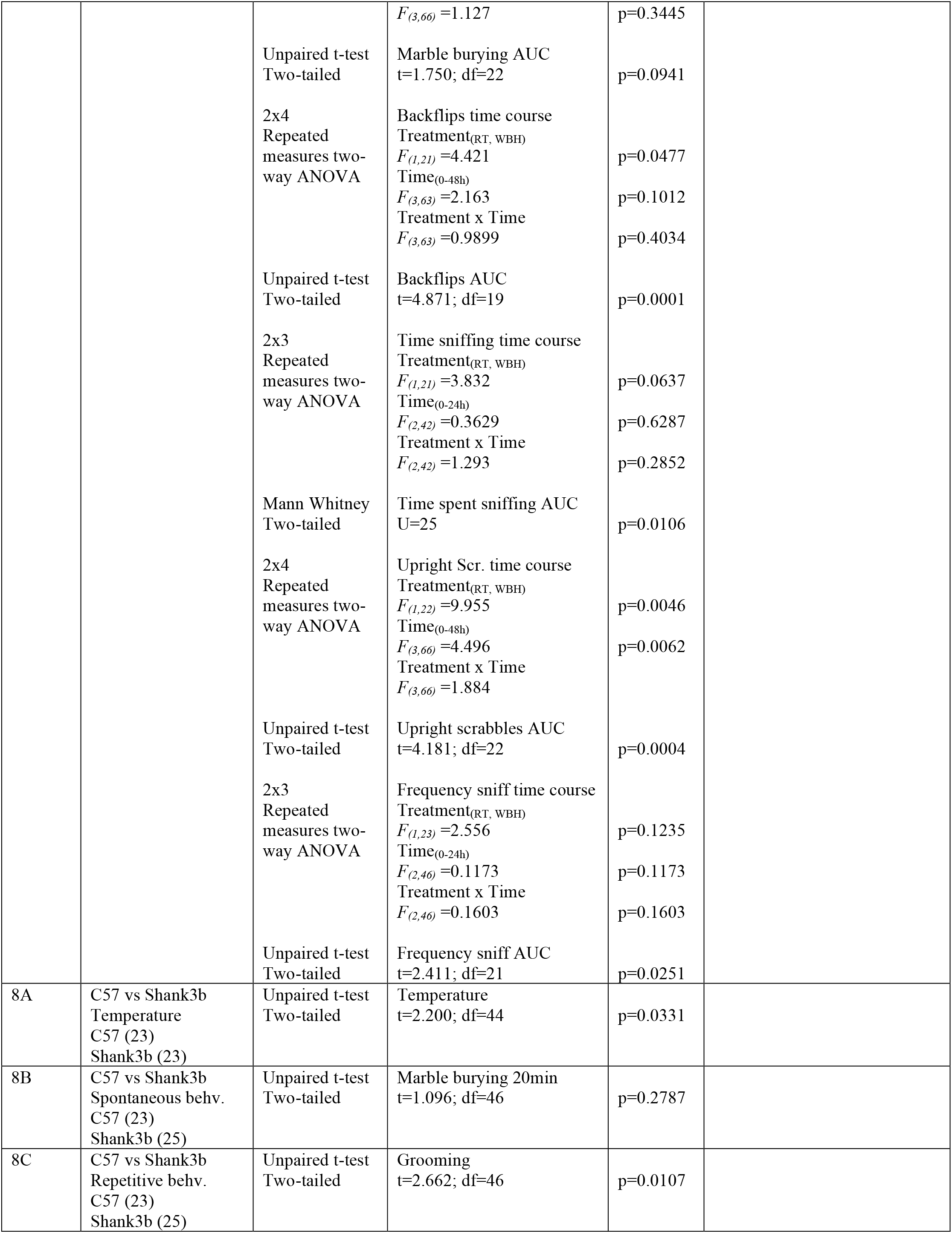
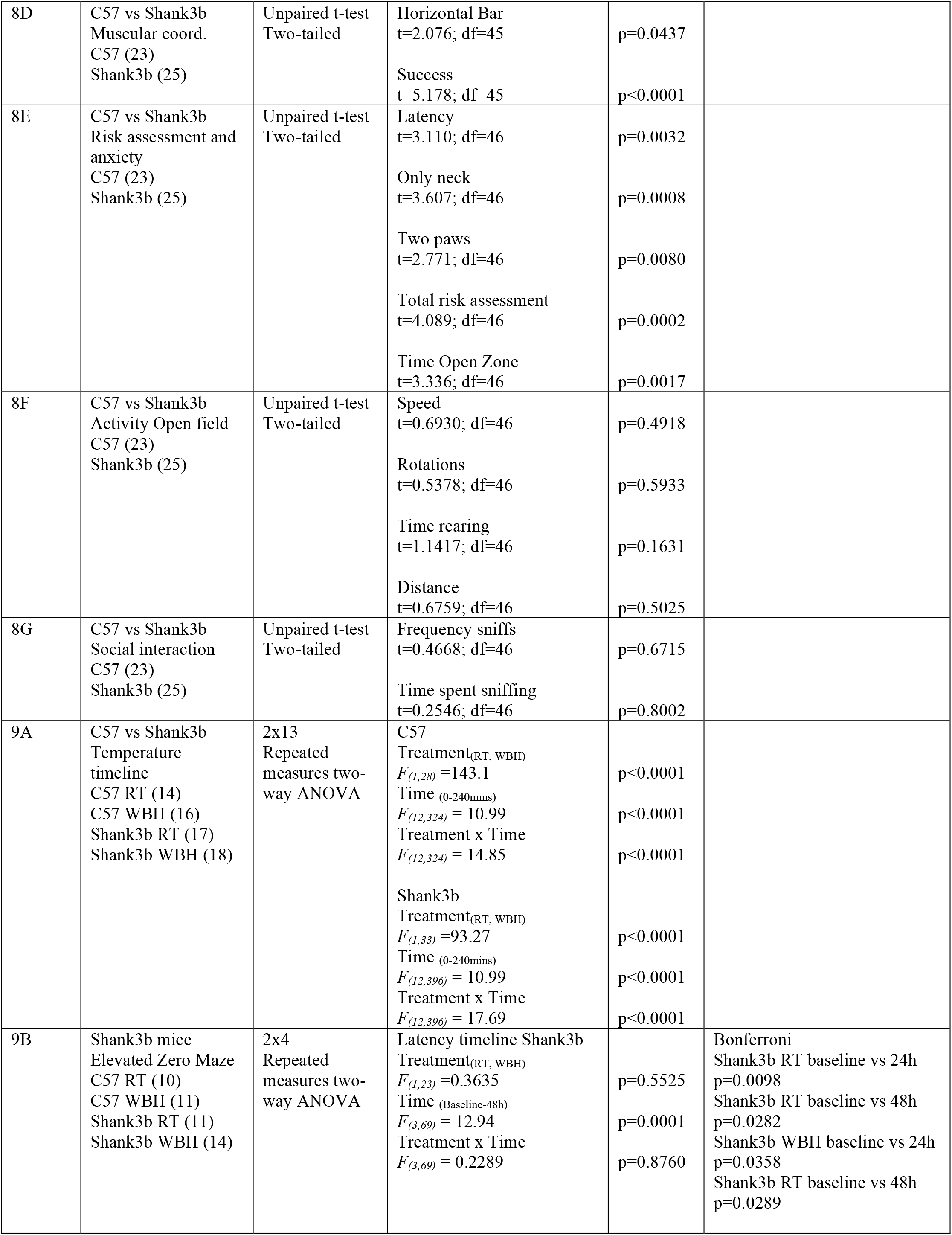
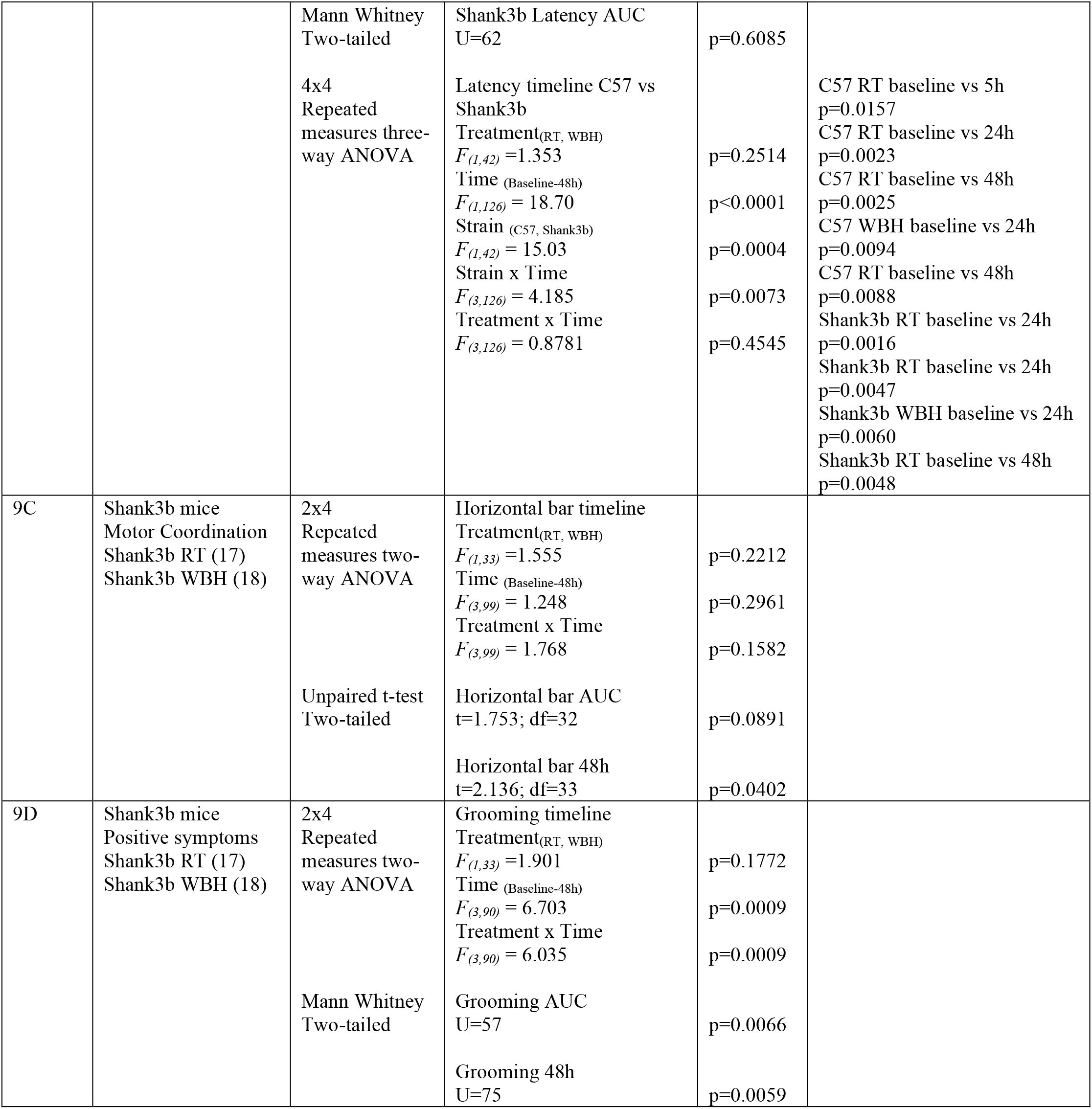
Summary of experimental design and statistical analyses used in this study

## Results

### Whole Body Hyperthermia protocol characterization

The experimental protocol is shown in Figure 1A. The Whole Body Hyperthermia (WBH) protocol induced an increase of body temperature reaching the target body temperature of 39.5 ± 0.5 °C (fever-like body temperature). Neither LPS-treated animals (250 μg/kg i.p.) nor RT animals showed a sustained increase in temperature from baseline, although LPS-treated remained at a significantly higher body temperature than RT mice (Figure 1B). Repeated measures two-way ANOVA showed a significant effect of treatment [F_2,69_ = 78.6, p<0.0001], of time [F_12,828_ = 40.61, p<0.0001] and an interaction between these two factors [F_24,828_ = 13.19, p <0.0001]. Subsequent Bonferroni’s multiple comparison test indicated a significant difference between RT and WBH, between WBH and LPS and between RT and LPS (all p<0.0001).

Body and brain temperature were also monitored simultaneously using subcutaneous transponders and brain-implanted thermocouples over the four hours duration of the WBH protocol (n=6 per group except n=5 for LPS) (Figure 1 C). Brain temperature was consistently significantly higher than body temperature, in all 3 treatment groups. There was a significant effect of body region when data were analysed by two-way ANOVA repeated measures in RT [F1,5=43.12; p=0.0012], LPS [F1,4=197.1; p=0.0001] and WBH [F1,5=65.27; p=0.0005] treated animals. The WBH protocol produces an acute elevation in brain temperature that persists as long as the WBH protocol continues, and returns to baseline temperature thereafter, just as it does in the body as a whole. When all three groups were pooled and analysed by linear regression, there was a statistically significant and strong positive correlation between body and brain temperature (r^2^=0.6742) and this persists even when body temperature is elevated, as in the case of the WBH animals.

For experiments in which subsequent behavioural assessment was necessary, the WBH protocol had to be refined by including an extra ‘step-down’ period (as per methods section) to allow animals to cool gradually in order to avoid rebound hypothermia (Wilkinson *et al*., 1988) (Figure 1A).

### Inflammatory markers induced by LPS and WBH

Since fever is typically underpinned by an acute inflammatory response, we sought to ascertain whether raised body temperature *per se* induced increased peripheral or central inflammation and whether this was comparable to inflammation induced by LPS. Plasma cytokine levels, assessed by ELISA, showed a clear induction of IL-1β, IL-6 and TNF-α in LPS-treated animals, but not in WBH animals (Figure 2A). Kruskal Wallis analysis revealed a significant effect of treatment on IL-1β, IL-6 and TNF-a levels [F_2,12_ = 9.750; p =0.004] with LPS significantly different to both RT and WBH in all cases, while WBH was not different to RT in any case. All statistical comparisons are shown in *table 2*. The results indicate that LPS significantly increases circulating inflammatory cytokines whereas WBH did not induce an inflammatory response.

**Figure 2.**
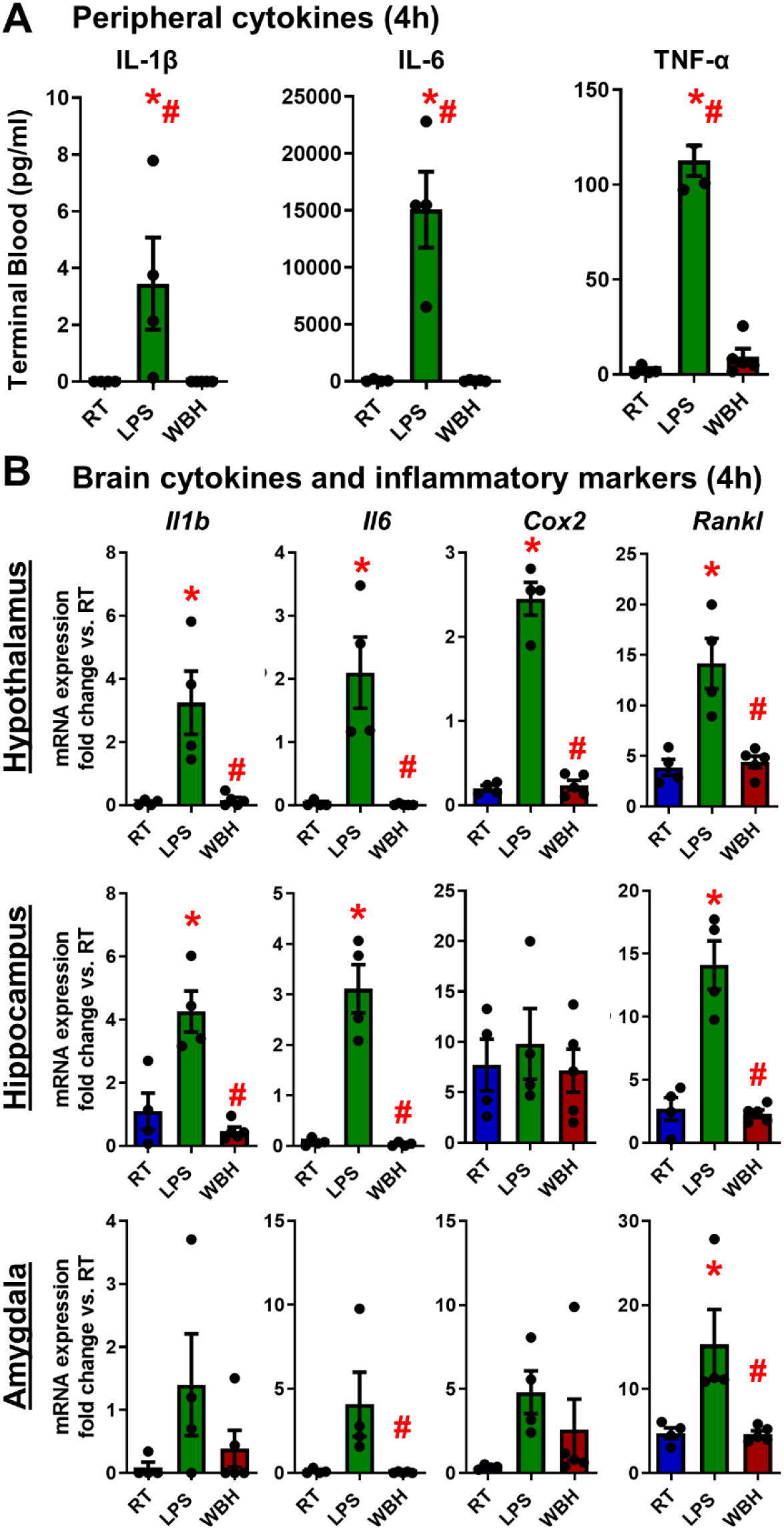
Inflammatory markers at 4h post Whole-Body Hyperthermia (WBH) or LPS. A) Peripheral blood cytokines measured at 4h after the WBH protocol: IL-1β, IL-6 and TNF-α levels (pg/ml). B) mRNA expression of brain cytokines and inflammatory markers in brain homogenates of hypothalamus, hippocampus, and amygdala. Data are shown as mean ± SEM, with each data point representing one animal (n=5 for WBH and n=4 for LPS, RT). Data have been analysed by parametric or non-parametric statistics as appropriate to the data distribution: Kruskal Wallis or One-way ANOVA followed by Mann-Whitney or Bonferroni’s tests p<0.05. (* LPS vs. RT; # LPS vs. WBH). Abbreviations: RT (room temperature); LPS (intraperitoneally-injected LPS, 250 µg/kg); WBH (whole-body hyperthermia).

Similar results were found in brain homogenates in three different regions in which *Il1b*, *Il6*, *Cox2* and *Rankl* mRNA levels were elevated by LPS treatment when compared to RT and WBH; however, WBH did not induce a brain inflammatory response and was not significantly different to RT controls on any gene in any region (Figure 2B). Conversely, LPS was typically significantly different both from RT control and WBH animals. *Rankl* was significantly induced by LPS in all regions examined, while *Il1b* and *Il6* were induced in the hypothalamus and hippocampus but were not significant in the amygdala owing to substantial variability in this region. *Cox2*, though robustly elevated in the hypothalamus, was not significantly induced in the hippocampus (high basal expression of *Cox2*) or in the amygdala (high variability). A summary of the statistical design and results is shown in table 2.

### cFos activation

Previous studies have shown that LPS treatment or environmental warmth (37°C) induce neuron activation following a differential brain-region pattern (Dallaporta *et al*., 2007; Tan *et al*., 2016). To characterize differential neuron activation patterns, cFos immunostaining was performed and analysed at 4 hours after WBH or LPS treatment. cFos showed different activation levels after WBH or LPS treatment in a brain region-dependent manner. WBH tended to induce cFos activation that was higher, and present in more brain regions, than LPS treatment did. Semiquantitative analysis of cFos positive cells (as described in the methods) is shown in figure 3. After examining cFOS activation in sections throughout the entire anterior-posterior axis, in an unbiased manner and with subsequent attention to regions previously suggested to be activated by systemic LPS or whole body hyperthermia, we observed three key patterns of activation with respect to whole body hyperthermia: i) equally induced by LPS and WBH: in paraventricular nucleus (PVN) (Bregma −0.46 to −0.94mm) and the ventromedial preoptic nucleus (VMPO) (Bregma 0.50 to −0.10mm); ii) more activated by the WBH protocol in comparison with RT or LPS treatment: the medial preoptic area (MPOA) (Bregma 0.76 to −0.58mm), the basolateral amygdala (BLA) (Bregma −0.58 to −3.16mm) and the dorsomedial hypothalamic nucleus (DMH) (Bregma −1.34 to −2.18mm); iii) exclusive activation pattern following WBH protocol: in the shell of the nucleus accumbens (ShAcc) (Bregma 1.94 to 0.74mm), the lateral septum (LS) (Bregma 1.94 to −0.10mm), the piriform cortex (PirCx) (Bregma 2.46 to −2.80mm) and the medial habenula (mHab) (Bregma −0.82 to −2.18mm).

**Figure 3.**
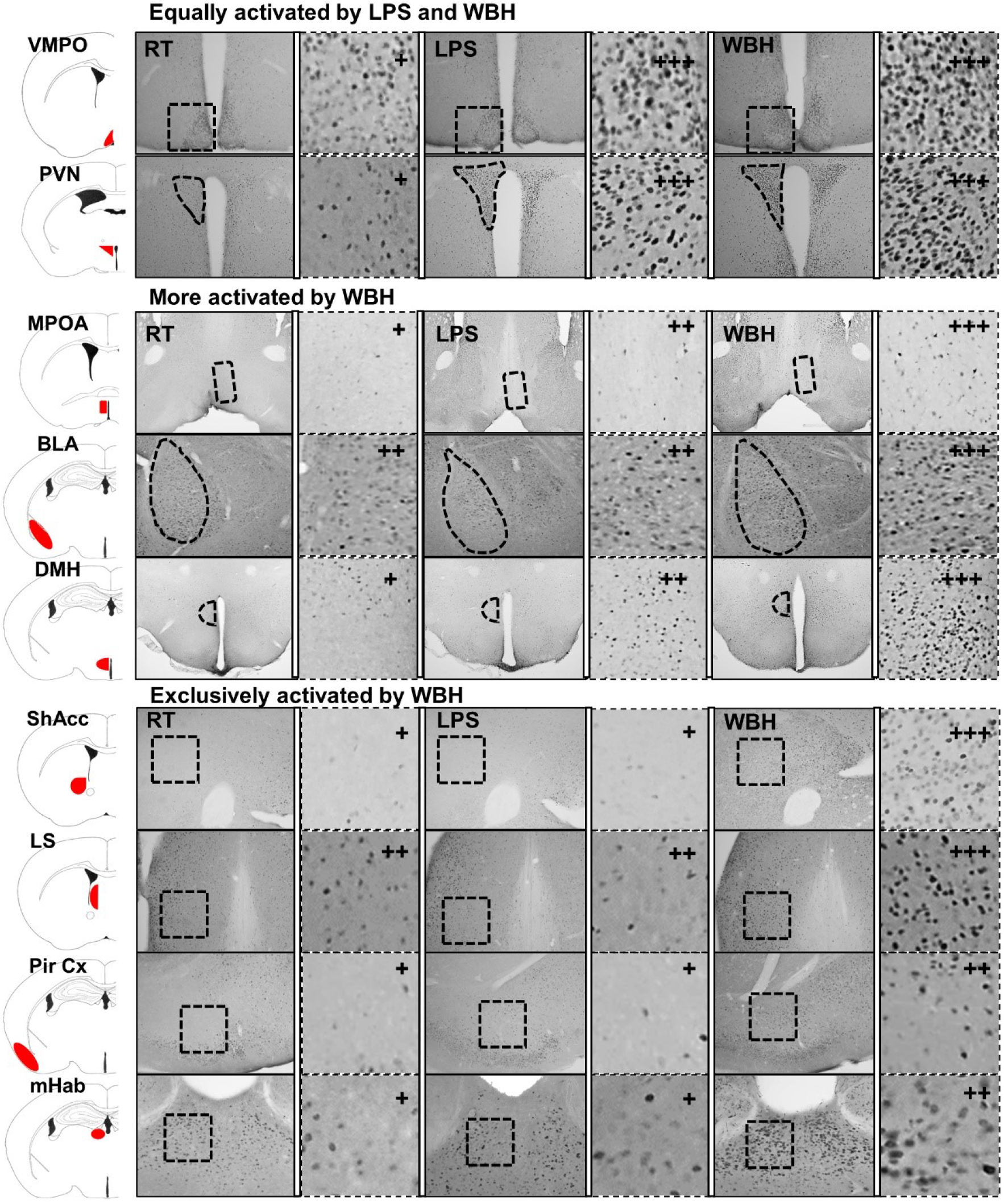
Differential patterns of labelling with the immediate early gene cFOS. Immunohistochemistry for cFos was performed in free-floating sections from brains of mice exposed to room temperature (RT), LPS (250 µg/kg) and whole body hyperthermia (WBH) (RT n=7; LPS n=8; WBH n=6). Semiquantitative analyses of neuronal activation induction: + (minimal activation); ++ (medium activation); +++ (high activation). Abbreviations: ventromedial preoptic nucleus (VMPO) (Bregma 0.50 to −0.10mm), paraventricular nucleus (PVN) (Bregma −0.46 to −0.94mm), medial preoptic area (MPOA) (Bregma 0.76 to −0.58mm); basolateral amygdala (BLA) (Bregma −0.58 to - 3.16mm), dorsomedial hypothalamic nucleus (DMH) (Bregma −1.34 to −2.18mm), nucleus accumbens (ShAcc) (Bregma 1.94 to 0.74mm), lateral septum (LS) (Bregma 1.94 to −0.10mm), piriform cortex (PirCx) (Bregma 2.46 to −2.80mm); medial habenula (mHab) (Bregma −0.82 to −2.18mm).

### Glucose levels, hypothalamic markers, and heat shock proteins

Significant metabolic changes are required to raise body temperature (Thorne *et al*., 2020) and also occur during the response to LPS (Kealy *et al*., 2020). Here, both LPS and WBH had a significant impact on plasma glucose levels when compared with control (RT) animals (Figure 4A). This was apparent in terminal bloods (4h post LPS or WBH) and, when using serial tail vein sample, could be shown to emerge as soon as 2 hours post-challenge (Figure 4B) and become statistically significantly different from RT in both groups by 4h (Main effect of treatment: F_2,23_ = 11.89, p=0.0003). The decrease was more profound in LPS-treated mice than in WBH, remaining significantly different to RT and WBH at 7h (p = 0.0037 and 0.0017 respectively) (Figure 4B).

**Figure 4.**
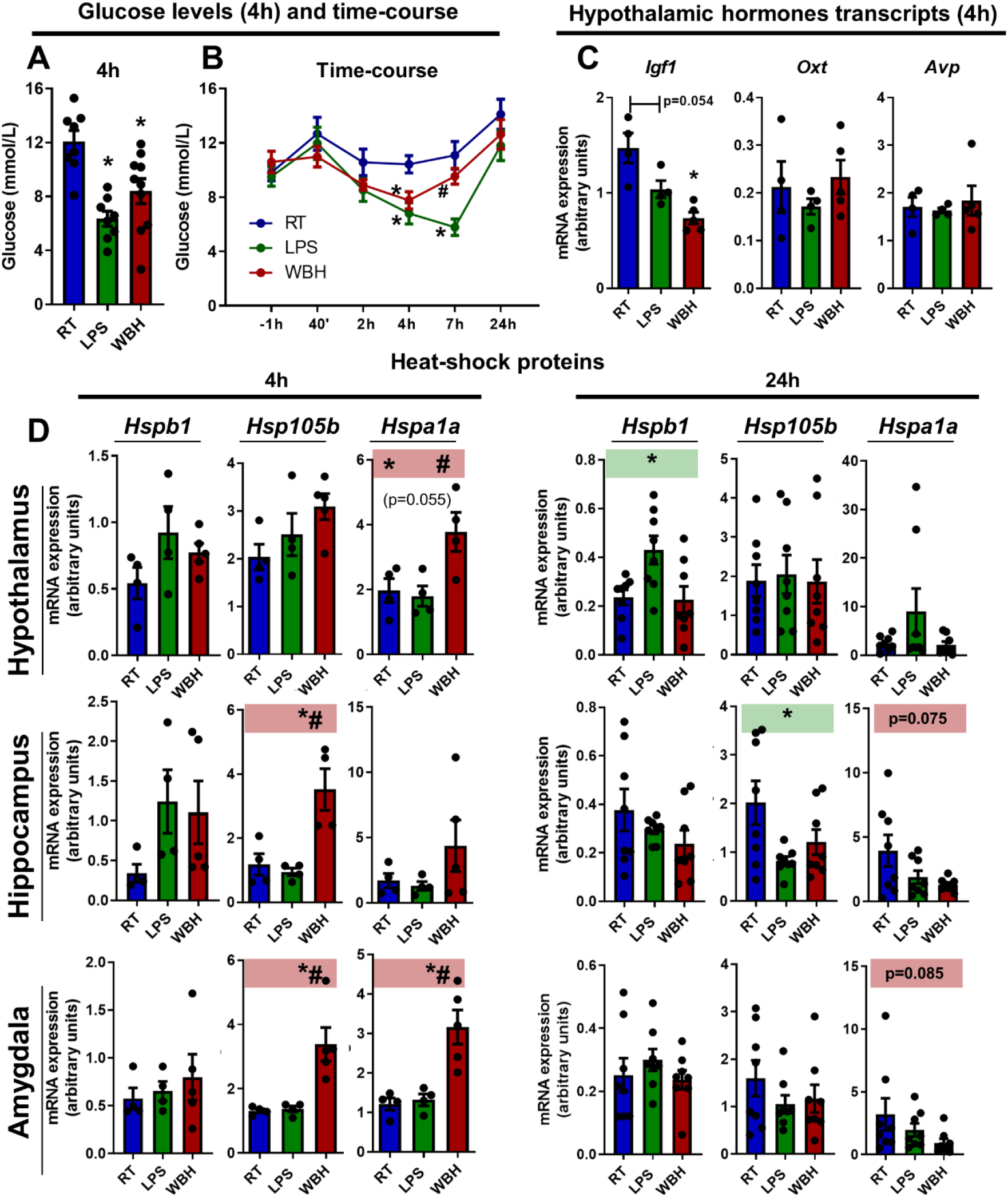
Glucose, hypothalamic hormone and heat shock protein mRNA levels after LPS and WBH protocols. A) Blood glucose levels (mmol/L) at 4 hours after LPS or WBH, shown as mean ± SEM (RT, LPS n=8, WBH n=10). B) Blood glucose levels (mmol/L) measured by serial tail vein sampling from 1 hour pre-to 24h post-LPS or WBH treatment (n=8). C) Level of expression of mRNA for hypothalamic hormones at 4 hours after LPS or WBH. D) Level of expression of mRNA for heat shock proteins in brain homogenates of hypothalamus, hippocampus, and amygdala at 4h (left, RT n=4; LPS n=4; WBH n=6) and 24h (right; n=8) after LPS or WBH treatment. Data are shown as mean ± SEM, with each data point representing one animal. Time course analysis was performed using two-way repeated measures ANOVA followed by Bonferroni post hoc analyses. All other analyses performed by one-way ANOVA followed Bonferroni’s test. * vs. RT; # vs. LPS. Abbreviations: RT, room temperature; LPS, intraperitoneally injected LPS, 250µg/kg; WBH, whole-body hyperthermia.

Effects of hyperthermia or LPS on mRNA levels were assessed for 3 hypothalamic-expressed hormones, all previously linked to ASD (Vahdatpour *et al*., 2016; Cataldo *et al*., 2018), 4 hours post-treatment (Figure 4C). One way ANOVA showed a significant effect of treatment on *Igf1* mRNA (F_2,10_ =12.78; p=0.0018), but not on *Oxt* or *Avp*. Post-hoc tests showed a significant decrease in *Igf1* only with WBH (p=0.0015).

Heat shock proteins (HSPs) are chaperone proteins that minimise protein denaturation and aggregation under heat shock (42-45 °C) and febrile-temperatures (38–41 °C), among other stressors (Di *et al*., 1997). *De novo* transcription of HSPs was analysed at 4 and 24 hours in brain homogenates of the hypothalamus, hippocampus, and amygdala (Figure 4D). Although quite variable, WBH did trigger increased expression of *Hspa1a* and *Hsp105B* in all three regions and these increases were significant in hippocampus and amygdala for *Hsp105b* (p=0.0038, p=0.0062, respectively) and in the hypothalamus and the amygdala for *Hspa1a* (p=0.0214, p=0.0017 respectively). None of these changes remained at 24 hours. Although there were some modest increases, neither WBH nor LPS significantly increased *Hspb1* and indeed LPS did not significantly increase any HSP at 4 hours. Conversely, *Hsp1b* was increased in the hypothalamus 24 hours after LPS treatment (p=0.0341) and *Hsp105b* was suppressed at 24 hours in the hippocampus (p=0.0332). See table 2 for full statistical descriptions.

### Effects of WBH and LPS in C57 mice

Bearing in mind the cellular and molecular changes observed in WBH and LPS mice, we assessed the impact of these stimuli on behaviour in normal C57 mice. Whole body hyperthermia did not significantly affect the normal behaviour of C57 mice whereas LPS induced a general decrease in activity as assessed in the open field test (Fig 5A). For the mean speed measurement, repeated measures two-way ANOVA showed a significant effect of treatment (F_2,29_ =4.121; p=0.0266) and time (F_2,58_ =9.898; p=0.0002) with Bonferroni post-hoc test showing the LPS-induced suppression of spontaneous behaviour at 5h compared with RT group (p=0.0206) and WBH (p=0.0080). Similar results were obtained for the number of rotations and reciprocal observations were made for the time spent immobile, showing that LPS significantly increased the time immobile at 5h when compared with the RT group (p<0.0001) and WBH (p<0.0001). Marble burying also demonstrated a suppression of this behaviour by LPS, while there were no effects of WBH. There was a significant effect of treatment (F_2,29_ =4.184; p=0.0253), which was driven by a significant effect of LPS at 5h, in comparison with RT group (p<0.0001) and WBH (p=0.0051). In the social interaction tests all animals showed suppression of social behavioural preference at 5 and 24 hours but there were no significant differences between RT, WBH and LPS (Figure 5C).

**Figure 5.**
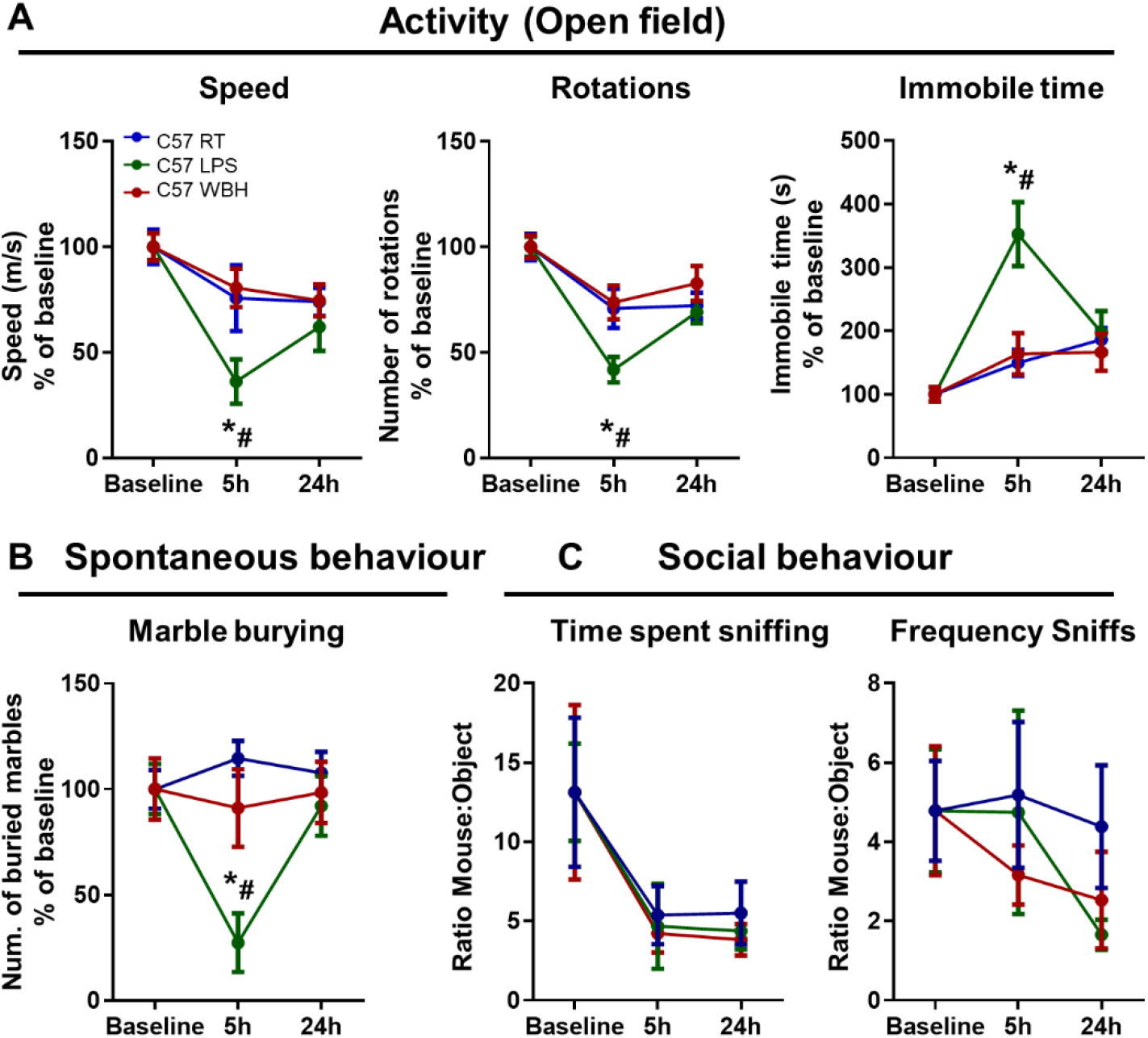
**Effects of WBH and LPS in C57 strain**. A) Description of the effects of WBH and LPS on the general activity using the open field test, showing the mean speed (m/s), number of rotations and immobility time (s) for 10 minutes. B) Characterization of the effects of WBH and LPS on social behaviour using the time of sniffing and the frequency of sniffs of a mouse in comparison with an inanimate object (ratio mouse:object) for 10 minutes. C) Analyses of WBH and LPS effects on the number of marbles buried within 20 minutes. Mean ± SEM (RT n=12; LPS n=8; WBH n=12). Repeated measures two-way ANOVA. * vs. RT and # vs. WBH by Bonferroni post-hoc tests. (RT: room temperature; LPS: intraperitoneally injected LPS, 250µg/kg; WBH: whole-body hyperthermia).

### Behavioural characterization of the C58 strain at baseline

The natural mutant C58 mouse was chosen as a suitable strain in which to assess the impact of WBH on autism-like features because they are reported to present several negative and positive symptoms/signs including modified spontaneous behaviours, stereotypy and highly characteristic repetitive movements like backflips and upright scrabbles, as well as decreased social interaction (Moy *et al*., 2008, 2014; Muehlmann *et al*., 2012; Blick *et al*., 2015; Whitehouse *et al*., 2017). In order to examine the impact of WBH on behavioral indices it was first necessary to replicate previously reported behavioral signs in C58 mice when compared to C57 mice. Basal body temperature was measured at 8 am, before assigning to treatment groups (Fig. 6A) and C58 mice were found to have a significantly lower basal body temperature than C57 mice (p=0.0001, unpaired two-tailed t-test, n=24 per strain). Burrowing of food pellets is a species-typical ‘natural’ behaviour in mice (Deacon, 2009; Jirkof, 2014) and was significantly increased in C58 mice both at 2 h (p=0.0273) and overnight (p=0.0101). Conversely, C58 mice showed significantly less marble-burying (p=0.0009), which can thus be described as a negative symptom in these mice. As in previous descriptions, we also observed a large number of backflips (p<0.0001) and upright scrabbles (p<0.0001) which were absent in C57 mice. These parameters may be regarded as “positive symptoms”. Hyperactivity has also been described in C58 mice and we used the open field test to show a significant increase in the speed (p<0.0001), number of rotations (p<0.0001) and travelled distance (p<0.0001) and a modest but significant decrease in the time spent rearing (p=0.0475) in C58 mice (Figure 6D, Unpaired t-tests). Finally, social behaviour was assessed using the social interaction test, indicating significant decreases in the frequency of (p=0.0142), and the time spent (p=0.0236), sniffing a novel mouse in comparison with an inanimate object (Figure 6E). There provide robust behavioural indices with which to assess the impact of WBH in proceeding experiments.

**Figure 6.**
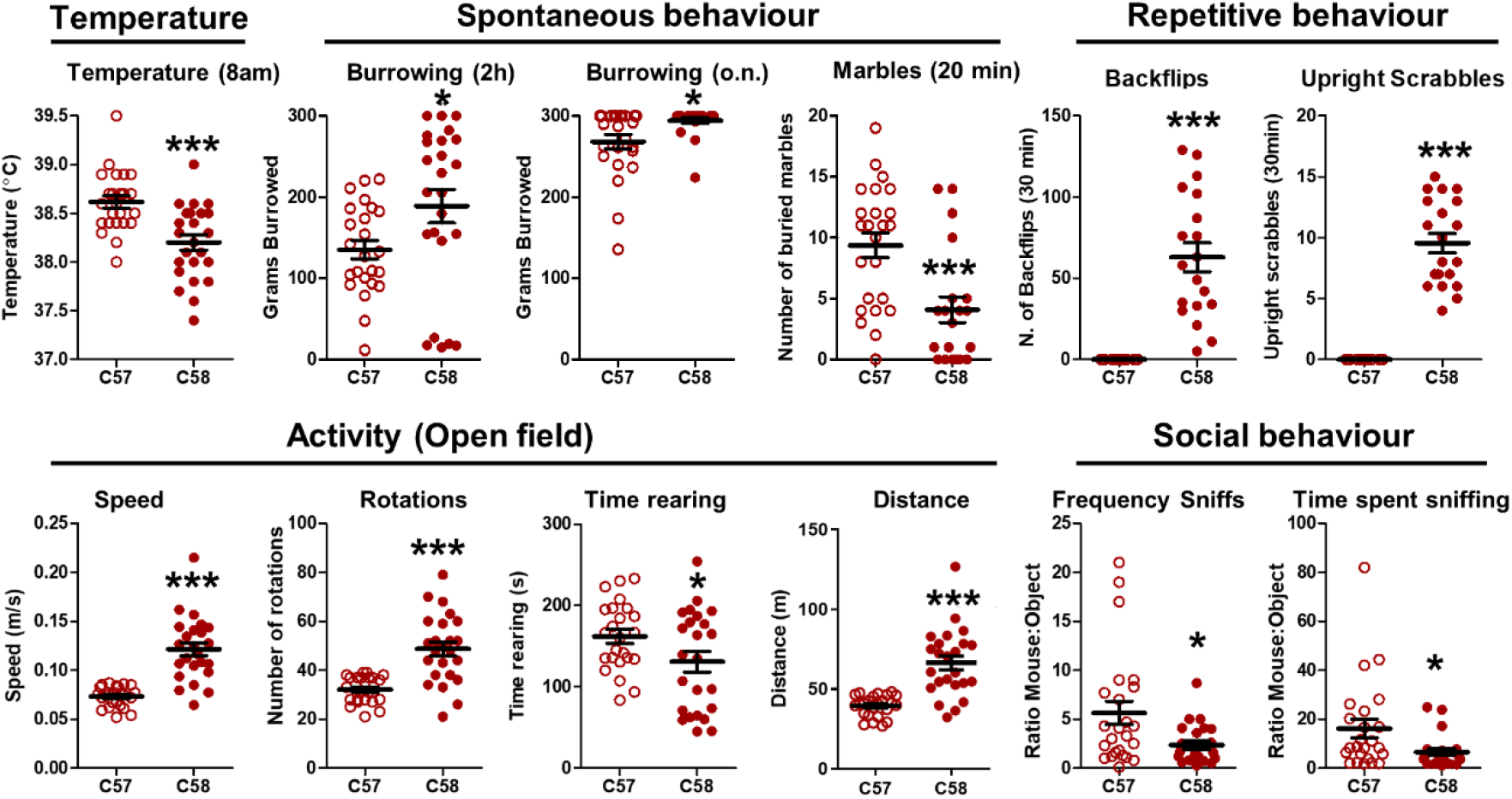
Characterization of basal behaviour in C58 mice. A) Basal body temperature at 8am, as measured by implanted transponder (°C). B) Spontaneous behaviour assessed by the amount of burrowed food pellets at 2h, and overnight, and by marble burying for 20 minutes. C) Description of the repetitive behaviours found in C58 strain: backflips and upright scrabbles for 30 minutes. D) Analyses of the general activity using the open field test, showing the mean speed (m/s), number of rotations, time rearing (s) and travelled distance (m) for 10 minutes. E) Social behaviour characterization using the frequency of sniffs and the time of sniffing of a mouse in comparison with an inanimate object (ratio mouse:object) for 10 minutes. Each animal is represented by a single data point and mean ± SEM are also indicated (n=24 for each strain). Unpaired two-tailed t-test, *,*** denote p<0.05, <0.001 for C58 vs C57 strain.

### Effects of WBH on C58 strain

The WBH protocol required adaptation to be used with C58 animals since they showed temperature increases above 40°C. In order to reach target body temperature of 39.5 ± 0.5 °C for the full 4 hours (p<0.0001, F_1,23_=139.2), the heating box was set at 33.5°C. WBH did not affect locomotor activity, as measured by mean speed or number of rotations in the open field test but did significantly reduce repetitive behaviours such as backflips and upright scrabbles (Figure 7C, left column). This improvement persisted when examined at 5, 24 and 48 hours. Repeated measures two-way ANOVA showed significant effects of treatment (F_1,21_ = 4.421; p=0.0477 and F_1,22_=9.955, p=0.0046) for backflips and upright scrabbles respectively. When analysed using area under the curve (AUC), WBH-treated C58 mice showed significantly lower levels of these repetitive behaviours (t-tests, p=0.0001 and 0.0004 for backflips and scrabbles respectively) indicating an improvement in key “positive symptoms”.

**Figure 7.**
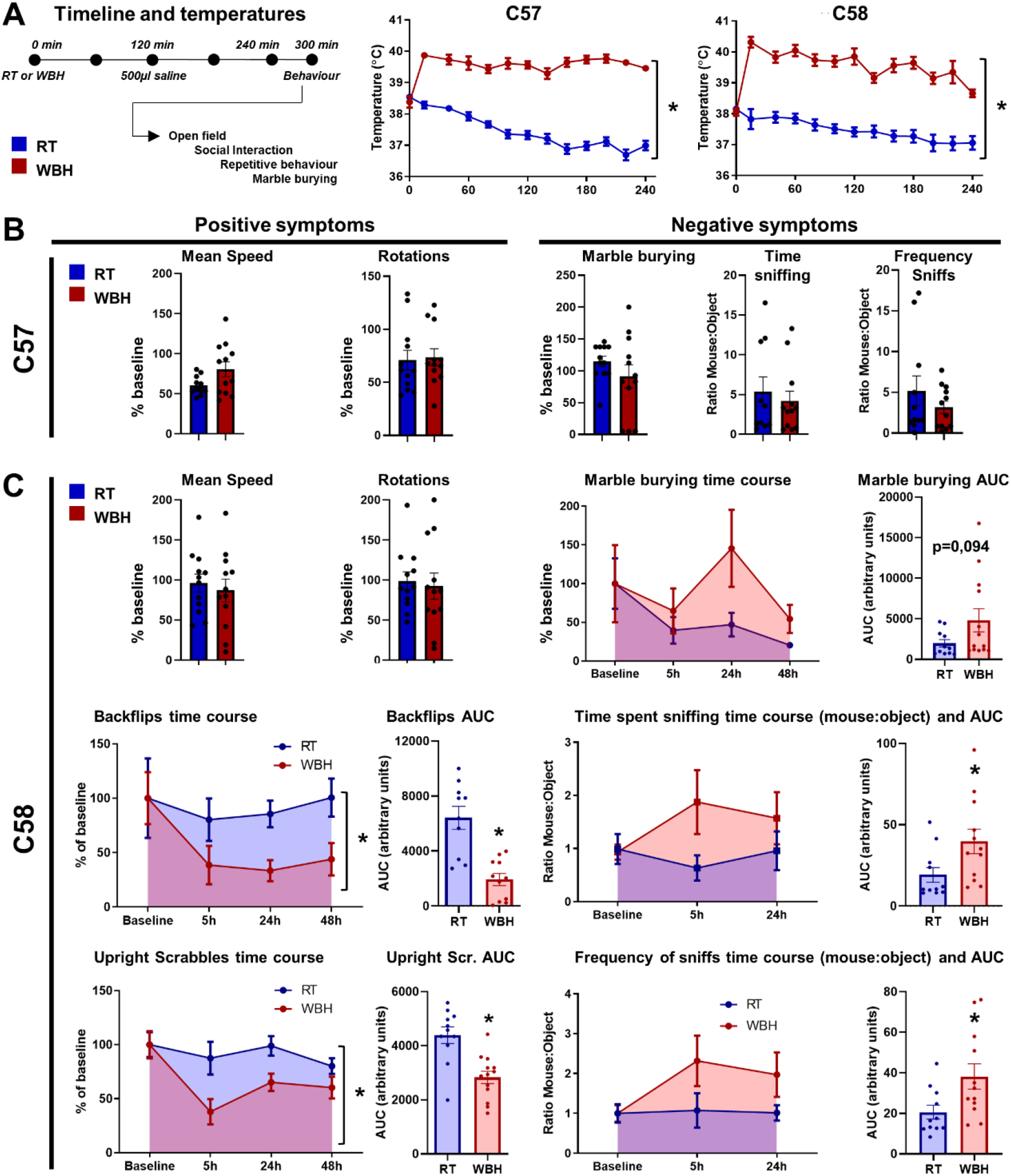
The impact of WBH on behavioural deficits in the C58 mice. A) Schematic timeline of the whole-body hyperthermia protocol and the behavioural tasks (left) and observed body temperature time course (°C) over the four hours duration of the protocol, measured by subcutaneous temperature transponders in C57 and C58 mice (right). The heating chamber was set to 33.5°C to reach the target body temperature of 39.5°C. B) Effects of WBH in C57 strain on positive symptoms assessed by the mean speed and number of rotations (left) and negative symptoms evaluated by the marble burying and the social interaction test (right). C) Effects of WBH in C58 strain in positive symptoms represented by mean speed, number of rotations and repetitive behaviours (left) and the effect of WBH on the negative symptoms measured by the marble burying and the social interaction test (right). Repeated measures two-way ANOVA. * denotes p<0.05 for WBH vs. RT by repeated measures two-way ANOVA, unpaired two-tailed t-test or Mann Whitney test.

Concerning “negative symptoms”, spontaneous behaviour in the marble burying test was somewhat increased by WBH, although this did not reach statistical significance: p=0.094 (Figure 7C, right column). WBH significantly improved social interaction producing an increase of the time-spent sniffing in WBH-treated C58 mice reflected by a significantly higher AUC (Mann-Whitney test, p=0.0106). This same result was found for the frequency of sniffs that showed a significantly higher AUC in the WBH group in comparison with the RT mice (student t-test, p=0.0251). Thus, WBH can improve both positive and negative symptoms in the C58 model, although not all behaviours were significantly mitigated.

### Shank3B- characterization in the basal state

In order to test whether fever range temperature was also beneficial in another ASD-relevant strain, we analysed Shank3B- mice. We chose heterozygotes since clinical autism-associated *Shank3* mutations are heterozygous. Similar to the C58 strain characterization, a battery of strain-specific behavioural tests were run on Shank3B- mice and normal controls, at baseline, in order to assess some autism-like positive and negative symptoms previously reported for homozygous Shank3B-/- mice (Peça *et al*., 2011; Mei *et al*., 2016; Balaan *et al*., 2019). Some, but not all, of those symptoms were observed here. Body temperature (at 8am), was significantly lower in Shank3B- mice in comparison with C57 mice (p=0.0331; Figure 8A). There was no significant difference between the 2 strains on marble burying (Thomas *et al*., 2009) (Figure 8B). Shank3B- mice are known to show characteristic excessive and repetitive grooming behaviour (Peça *et al*., 2011; Mei *et al*., 2016; Balaan *et al*., 2019) which we also observed here, as increased time spent grooming (p=0.0107) in Shank3B- mice (Figure 8C). Muscle strength and coordination were assessed using the horizontal bar test (Deacon, 2013a) and Shank3B- mice spent a significantly longer amount of time to reach the lateral platform (p=0.0437) and presented a significantly lower rate of success to complete the horizontal bar test (p<0.0001) (Figure 8D). The elevated zero maze (EZM) was used to assess anxiety-like behaviours and the trade-off between motivation to explore and tendency to remain in the less anxiogenic closed area (Deacon, 2013b). At basal levels, Shank3B- mice showed significantly higher latency to go to the open zone of the EZM (unpaired t test p=0.0032), spent less time in the open zone of the EZM (p=0.0017), (Figure 8E) and showed a higher frequency of risk assessment (see table 1 for full statistical analysis). General and exploratory activity were not significantly different between strains (Figure 8F). We observed no significant differences in the social interaction tests, either by time spent, or frequency of sniffing a novel mouse (Figure 8G). This battery identifies several useful measures with which to assess the effect of WBH on behavioral features of the Shank3B- strain.

**Figure 8.**
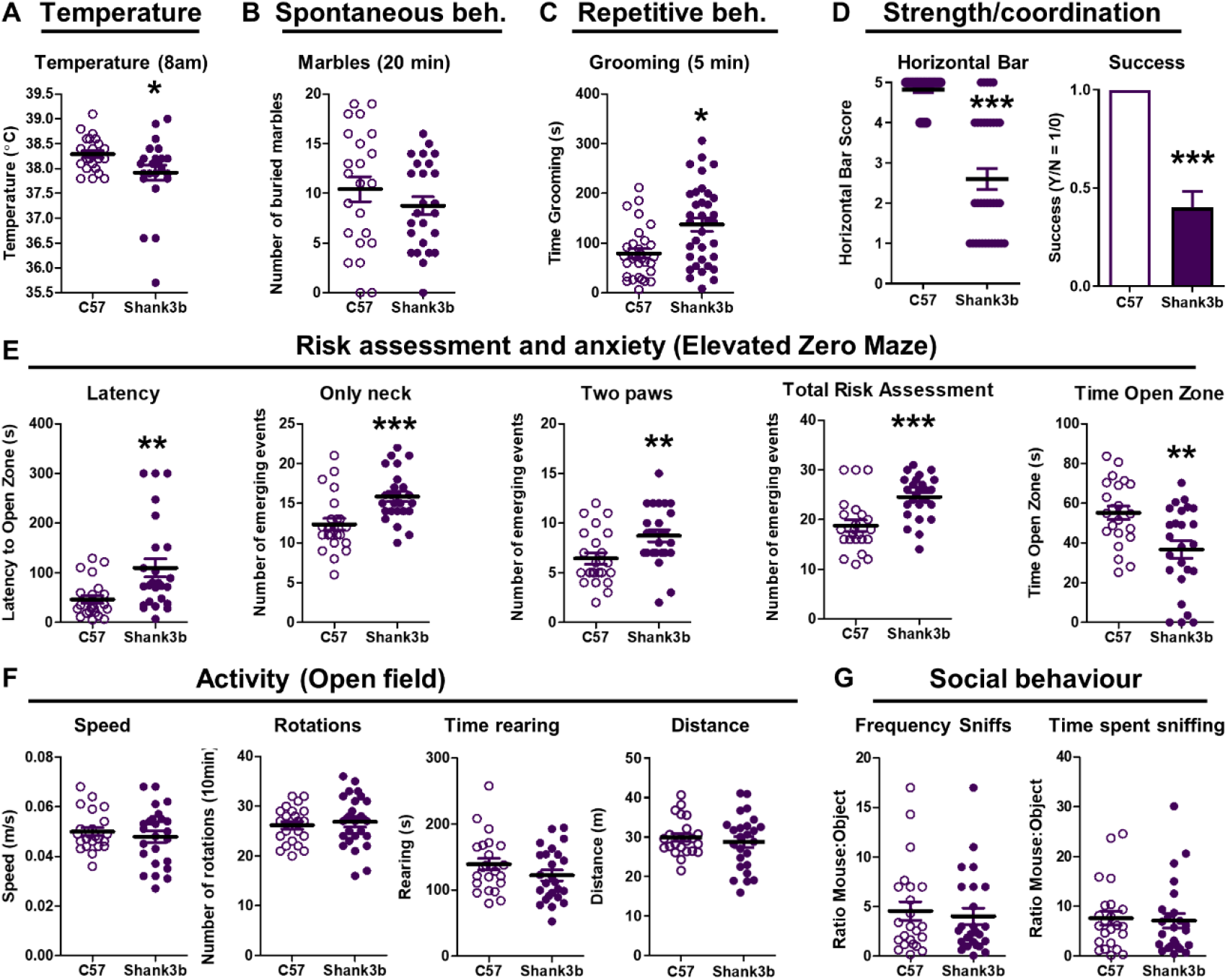
Characterization of Shank3B- mice in basal state. A) Body temperature, basal state at 8am, as measured by implanted transponder (°C). B) Spontaneous behaviour as measured by marbles buried in 20 minutes. C) Description of the repetitive behaviours found in Shank3B- strain: grooming test for 5 minutes. D) Assessment of the motor coordination and muscular strength on the horizontal bar test for 1 minute. Animals score 1 if they fell off within 10 seconds, score 2 if they held on for 11-59 seconds, score 3 if they held on for 60 seconds or reached the safe platform in 60 seconds, score 4 if they reached the safe platform within 30 seconds and score 5 if they reached the platform within 10 seconds. E) Analyses of risk assessment and anxiety-like behaviours using the elevated zero maze (EZM) test and monitoring the latency time to go to the open zone of the maze, the number of events of risk assessment (neck, two-paws, and total events) and the time spent in the open zone of the EZM during 5 minutes F) General and exploratory activity as assessed in the open field test, showing the mean speed (m/s), number of rotations, time rearing (s) and travelled distance (m) for 10 minutes. G) Social behaviour characterization using the frequency of sniffs and the time spent sniffing of a mouse in comparison with an inanimate object (ratio mouse:object) for 10 minutes. All data are expressed as mean ± SEM, n=23 per group. * vs C57 strain by unpaired two-tailed t-test, p<0.05.

### Effects of WBH on Shank3B- strain

Using the WBH protocol and behavioural follow-up (shown in Figure 9A) Shank3B- animals and C57 control mice reached the target temperature of 39.5 ± 0.5 °C and maintained this during the four hours of the protocol (significant effect of WBH treatment in C57 and Shank3B- mice (F_1,27_ =143.1, p <0.0001 and F_1, 33_=93.27, p <0.0001 respectively). Repetitive and excessive grooming is one of the specific alterations of Shank3B- mice. It is an abnormal behaviour that sometimes leads to skin lesions and scarring. Grooming was significantly improved after WBH (Figure 9B), reflected in a significantly lower AUC when assessed repeatedly over 48h (Mann Whitney t-test on AUC, p=0.0170) and this effect was maintained up to 48h after WBH (Mann Whitney t-test, p=0.0059). WBH did not significantly improve muscle strength/coordination on the horizontal bar. When measured repeatedly over 48 hours, performance of WBH animals decreased at 5 and 24 hours before returning to normal at 48 hours. The AUC was slightly lower in the WBH group, but this did not reach the statistical significance (p=0.080; figure 9C). Although Shank3B- showed higher latency to exit the closed area in EZM (as per figure 8E), this task proved ill-suited to test/re-test over 48 hours since the maze became less anxiogenic on repeated exposures in all mice (Figure 9D). As such it was not possible to demonstrate any effect of WBH on this measure of anxiety.

**Figure 9.**
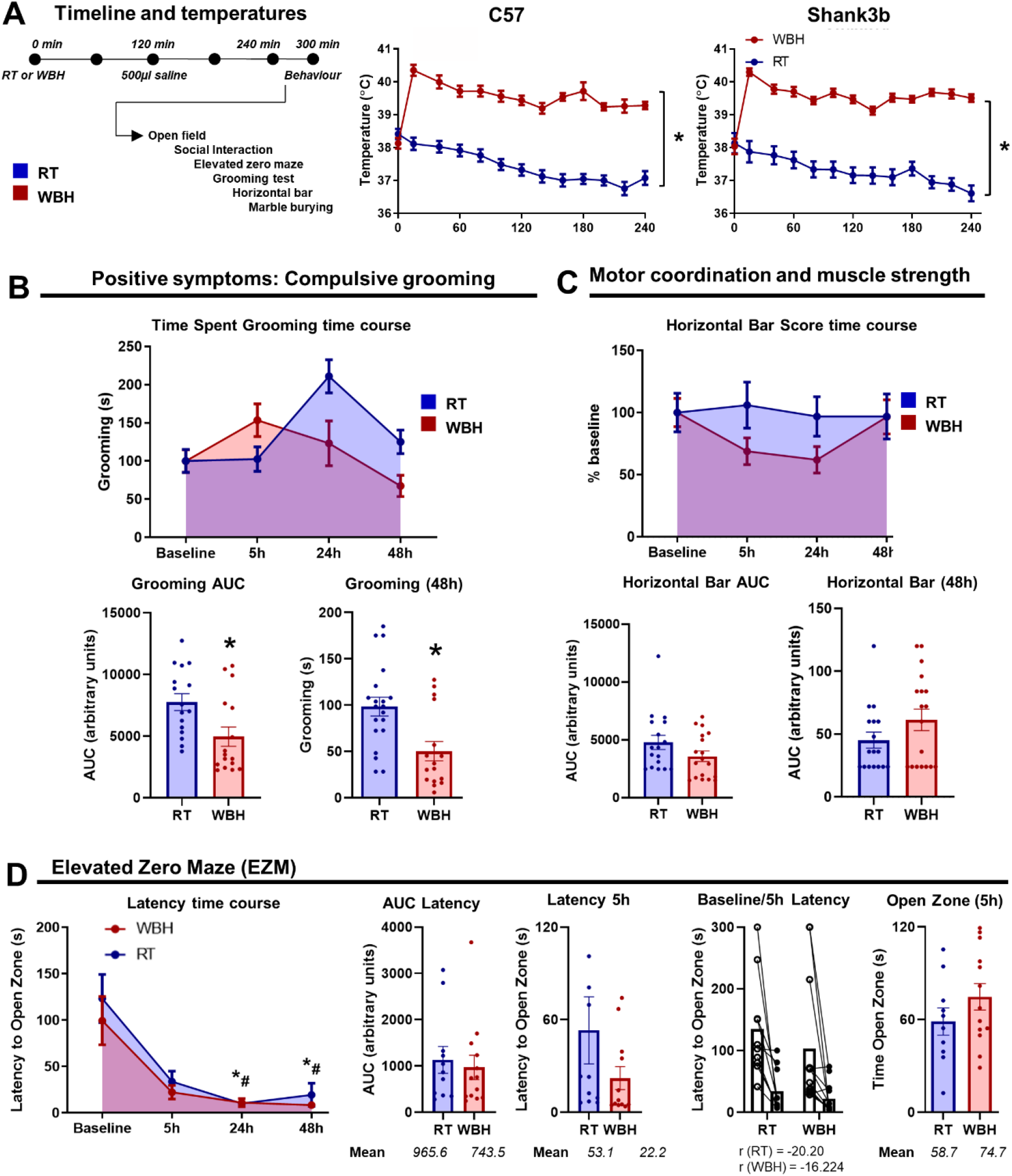
Effect of WBH on behavior in Shank3B- mice. A) Schematic timeline of the WBH protocol and behavioural tasks undertaken (left) and the impact of that protocol on body temperature (°C) over the five hours duration of the protocol, measured by subcutaneous temperature transponders in C57 and Shank3b- mice (right). B) Effects of WBH on time spent grooming in Shank3B- mice at 5, 24 and 48 hours (up) with AUC calculation for the entire period (down), and performance at 48h. C) Effect of WBH on motor coordination and muscular strength in Shank3B- and C57 mice, as assessed by the horizontal bar test. Performance of the horizontal bar based on the percentage of baseline scoring (up) and AUC (down) for measures over 48 hours post-heating, as well as performance at 48h are shown. D) Effect of WBH on latency to enter the open zone on the elevated zero maze in Shank3B-, shown across time and by AUC. Data are presented as mean ± SEM (Shank3b RT n=17; Shank3b WBH n=18) and are analysed by repeated measures two-way ANOVA, p<0.05 vs. RT group denoted by *; or by unpaired two-tailed t-test or Mann Whitney test, p<0.05 vs. RT group denoted by *.

## Discussion

The present study experimentally dissociates the inflammatory and hyperthermic components of fever to show that raising body temperature to 39.5 ± 0.5 °C for 4 hours, in the absence of inflammation, can improve symptoms in two distinct mouse models of ASD.

### ***D***issociation of hyperthermia from inflammation

We deliberately dissociated hyperthermia from inflammation in the current study since the sickness behavioural changes induced by acute systemic inflammation, including lethargy, a general suppression of spontaneous activity (Kealy *et al*., 2020), might produce apparent improvements in ‘positive symptoms’ (i.e. behaviours that show higher frequency in the ASD model of choice). Here, LPS-treated animals also showed the expected suppression of behaviours more generally and did not produce any lasting mitigation of either positive or negative symptoms as defined herein. In the most recent human study of this phenomenon very few children were reported to benefit from febrile episodes, but the examination of individual features revealed effects consistent with sickness behaviour responses (increased sleep but negative effects on activity, feeding and happiness). This is consistent with the idea that acute illness per se is unlikely to provide improvements for individuals with ASD. Whether elevated body temperature alone may provide benefits is a distinct question. Heat and inflammation were fully dissociable in our experimental manipulations. Only hyperthermia, induced by WBH, significantly elevated body and brain temperature (Figure 1) while only LPS (250 µg/Kg) induced pro-inflammatory cytokines (IL-1β, IL-6 and TNF-α) in peripheral blood and expression of *Il1b*, *Il6*, *Cox2* and *Rankl* mRNA in the hypothalamus, hippocampus, and amygdala (Figure 2). The WBH protocol, adapted from previously published WBH experiments (Ostberg *et al*., 2000; Evans *et al*., 2015) increased body temperature to a target temperature of approximately 39.5°C that was maintained during the four hours of the protocol. LPS treatment did modestly elevate body temperature with respect to RT controls, but this did not reach the fever range and indeed was not elevated from that observable at 8 am (immediately following the dark phase). Thus, only WBH protocol induced fever-range temperature.

It has recently been shown that acute inflammation induced by LPS, triggering expression of IL-17a, improved social behaviour in a maternal immune activation (MIA) model of ASD (Reed *et al*., 2020). This suggests that inflammation occurring during a fever response could contribute to improved performance in some behaviours. Comparison with the current study is necessary. Firstly, in the Reed study, improvements in social behaviour occurred specifically in those individuals whose neurodevelopmental abnormalities came about through *in utero* inflammation and did not occur in monogenic models including the Shank3B- model used in the current study. This is likely to be significant since there are clear data that responses to acute inflammatory stimuli are strongly influenced by prior inflammatory exposures, including those occurring *in utero* (Meyer, 2014; Neher & Cunningham, 2019). The LPS dose used by Reed et al., was lower than that used here (50 µg/Kg vs 250 µg/Kg) and this produced a more modest suppression of activity than we observed, which may facilitate a maintenance of social activity. The higher dose used in our study may have produced lethargy that was incompatible with observing improved performance on some motivated tasks (Figure 5) and which is consistent with illness providing limited benefit for ASD patients (Byrne *et al*., 2022). The current study is focussed on elevated temperature in the absence of acute illness. Reed et al., exclude an effect of fever *per se* in their LPS/MIA studies but we propose that this is premature since neither inflammatory-mediated, nor DREADD (designer receptors exclusively activated by designer drugs)-mediated increases in body temperature reached the fever range (always remaining <37 °C). In the current study we observed improvements in behaviour in both C58 and Shank3B- models after animals reached body temperatures of 39-40 °C. Using a novel thermocouple, we also showed that the brain temperature increased temporally in synchronisation with, and proportionately to, body temperature. In general, brain temperature was approximately 1.5°C higher than body temperature independently of the treatment (RT, LPS or WBH). Whether this elevation of brain temperature is necessary for the behavioural effects observed now requires investigation.

### WBH alleviates strain-dependent features in C58 and Shank3B- mice

It is significant that WBH significantly mitigated some behavioural and physiological features in these two widely used models of ASD, although not improving all behavioural features, and not completely reversing those that were improved. Given that ASD comprise conditions that are largely genetically determined and developmental in nature, any environmental change that can improve function deserves interrogation.

In the two strains examined here, the WBH protocol significantly decreased the key repetitive behaviours associated with those specific models. That is, backflipping and upright scrabbles in C58 mice and excessive grooming in Shank3B- mice. Both backflipping in C58 mice and excessive grooming were reduced by more than 50% and this effect persisted at 48 hours, two days after animals had returned to normal body temperature. Given the observed improvements in ASD individuals with more severe repetitive behaviours (Grzadzinski *et al*., 2018), the current findings may be significant. However, the reduction of hyperactive (or positive) behaviours is not, of itself, sufficient to demonstrate improvement in ASD-related phenotypes. This is because in the initial fever studies (Curran *et al*., 2007) individuals with fever typically also experienced lethargy and this might have contributed to apparent improvements in repetitive behaviours. Our studies with LPS (figure 5) clearly showed suppression of spontaneous behaviour during acute inflammation, making this stimulus unsuitable for assessment of changes in ASD-related behaviours. Given the scope for non-specific suppression of general activity, it was also important to show that WBH could also trigger improvements in “negative symptoms”, i.e. behaviours that are suppressed in the C58 and Shank3B- strains also showed improvements in WBH treated animals. There was a significant improvement in social behaviour in C58 mice.

These data illustrate that there may be significant behavioural benefits that outlast raised body temperature and stimulates important questions about whether there may be significant changes in synaptic plasticity arising from these fever range temperatures. Body and brain temperature have been shown to fluctuate across the circadian clock, to be approximately 1.5°C higher than arterial blood and to change with environment stimulation (Kiyatkin et al., 2002). Increased brain temperature in the dark phase, and upon initiation of activity, has recently been shown to influence hippocampal neurophysiological activity (Petersen et al., 2022) and raising the temperature of hippocampal slices has been shown to increase excitatory post-synaptic potentials (Schiff & Somjen, 1985). Experimental and modelling studies of optogenetic approaches indicate that brain tissue proximal to fibre optic stimulation (10 µm) can rise by as much as 1.5 – 2°C and modelling of this temperature change predicted increased firing frequency of interneurons, expedited gamma oscillation development and increased synchrony of neuronal firing (Peixoto et al., 2020). AMPA receptors show more rapid and larger peak amplitudes at higher temperatures, underpinned by more rapid post-synaptic AMPA receptor kinetics (Postlethwaite et al., 2007). How these changes affect synaptic summation in a complex network is more difficult to model and predictions for behaviour are unclear. Nonetheless, in the current study, raising body and brain temperature by approximately 1.5-2°C improved several behavioural indices in two ASD models.

We used Shank3B- heterozygotes to examine the impact of whole-body hyperthermia on mice with this ASD-relevant mutant. Most studies using Shank3B- mutants have compared null mutants to wildtype littermate controls (Peça et al., 2011; Wang et al., 2011; Mei et al., 2016; Dhamne et al., 2017). However, ASD cases, particularly those with Phelan-McDermid syndrome (Phelan & McDermid, 2012) are largely haploinsufficient rather than null (Betancur & Buxbaum, 2013) and deletion or loss-of-function mutations of one copy of SHANK3 may account for 0.5–2.0% of ASD and intellectual disability (Leblond et al., 2014). Thus haploinsufficiency of Shank3B- is sufficient to convey a high risk for ASD in humans and is sufficient for synaptic deficits and altered social interaction and communication in mice (Bozdagi et al., 2010). Shank3+/ΔC heterozygotes (C-terminal, exon 21-deleted Shank3), which exhibit reduced full-length Shank3 expression, also show significant repetitive behaviours and social deficits (Duffney et al., 2015; Qin et al., 2018). Here, we observed that Shank3B- phenotypes were largely still evident in heterozygous Shank3B- mice, although we did not observe any clear social impairments. Of the clear behavioural phenotypes observed, some key deficits were mitigated by fever range temperature. The basis of this temperature-induced improvement has not been determined here, so we can only speculate on what the mechanisms may be, but a consideration of Shank3B function maybe instructive.

Shank3B is a scaffolding protein in post-synaptic dendrites at glutamatergic synapses and interacts with PSD95 to form a post-synaptic complex which also facilitates interaction with NMDA, AMPA and metabotropic glutamate receptors. Dendritic structure and function is markedly impaired in Shank3B mutants. When Shank3B interaction with AMPA receptors was prevented, through deletion of Shank3 ankyrin repeat domains, excitatory AMPA-mediated transmission and hippocampal LTP were reduced in a manner reversed by AMPA receptor stimulation (Bozdagi et al., 2010). Shank3B-/- mice are highly resistant to PTZ-induced seizures and show enhanced gamma band oscillations, indicative of enhanced inhibitory tone (Dhamne et al., 2017). It is tempting to speculate that increased temperature is enhancing glutamatergic activity and normalising excitatory/inhibitory balance.

Interestingly there are also changes in dendritic structure in the inbred strain C58/J. These mice show several ASD-like behaviours, but the genetic basis of neuronal circuit dysfunction is unclear. However single nucleotide polymorphisms in genes for dendritic proteins including MAP1A (Microtubule-Associated Protein 1A), GRM7 (Metabotropic Glutamate Receptor, 7), ANKRD11 (Ankyrin Repeat Domain 11) have been reported, suggestive of impaired dendritic function at glutamatergic synapses (Barón-Mendoza *et al*., 2021). Both repetitive and social interaction deficits in C58 mice were mitigated by mGluR5 negative allosteric modulation (Silverman *et al*., 2012). Thus, there is a basis for the idea, although speculative, that modulating glutamatergic transmission may mitigate behavioural abnormalities in these two strains and that increasing temperature may facilitate this.

### Dissociable pathways of cellular and molecular activation

Although we have not described the cellular or molecular mechanisms for temperature-driven mitigations of behavioural deficits in the C58 and Shank3B- mice, a short examination of those cellular and molecular changes that differentiate WBH and LPS may be useful.

Using cFos labelling, primary and downstream regions previously described in a ‘warm sensitive circuit’ that responds to elevations in internal or external temperature (Tan *et al*., 2016), were clearly activated: anterior and preoptic areas of the hypothalamus mediate thermoregulatory homeostasis and lateral septum, medial habenula, BNST and PVN also participate in this circuitry.

As well as the BNST, PVN and VMPO, activated 4h after LPS or WBH, cFOS labelling identified several nuclei preferentially (MS, MPOA, BLA) or exclusively (ShAcc, LS, PirCx, mHab) activated by fever range temperature (Fig 3). Many of these regions are part of the extended amygdala circuit, involved in anxiety and fear (Shackman & Fox, 2016). The ShAcc activation observed, appeared particularly selective for WBH and the ShAcc is known to be subject to excitatory enervation from the BLA, which increases reward seeking and connectivity between these regions, and the BNST, is thought to facilitate behavioral responding to salient cues (Stamatakis *et al*., 2014). The PVN, amygdala and hippocampus are implicated in emotional and memory contributions to social interaction (Young *et al*., 2006; Wang *et al*., 2014; Tzakis & Holahan, 2019) and the mHab plays an important role in social interaction and social play behaviours (van Kerkhof *et al*., 2013). Interestingly there is also evidence that warm temperature can activate the interfascicular dorsal raphe (DRI) through the spinoparabrachial pathway, leading to activation of LS, NAc, mHab, dorsal hypothalamus and other regions (Hale et al., 2013) and can alleviate some symptoms of depression in those with major depression (Hale *et al*., 2013; Hanusch *et al*., 2013; Janssen *et al*., 2016). The mechanisms of those effects are entirely unclear, but it will be of interest to use optogenetic or DREADD (designer receptors exclusively activated by designer drugs) approaches to experimentally activate selected brain regions identified herein with a view to simulating the positive effects of fever on ASD-related behavioural abnormalities.

Hypoglycemia developed by 4 hours under both LPS and WBH treatments (Fig 4 A-B) and this is shown to trigger reduced spontaneous activity (Kealy *et al*., 2020). However glucose levels began to return to baseline levels shortly after animals were removed from the heating chamber and are unlikely to be the primary driver of the behavioural improvements seen with WBH. However, acute hypoglycemia drives elevated blood ketones and ketone diets have been suggested to be beneficial in some ASD cases (Li *et al*., 2021). Given the multiple bioenergetic changes that may occur during elevated ketone levels, it may be informative to assess ketogenesis during WBH and to assess general impacts on brain bioenergetic function.

Heat Shock Proteins are traditionally thought to only be involved in heat shock (42-45 °C) but can be induced by febrile-temperatures (38 – 41 °C), as seen here in the hypothalamus, hippocampus and the amygdala. The fold induction was modest and not occurring with all assessed genes (*Hspa1a, Hsp105B*). It remains to be seen whether these changes in transcription translate into elevated protein levels. Given HSPs role as chaperone proteins, minimising protein denaturation during stress (Di *et al*., 1997), it is not intuitive that this chaperone function would, of itself, offer a benefit to C58 and Shank3B- mice unless ongoing deficits in protein folding or conformation exist, however protein-protein interactions are central to many ASD-association mutations (Chen *et al*., 2018) and misfolding has been described in Neuroligin mutants (De Jaco *et al*., 2010). In principle chaperones such as HSPs might help stabilise molecular scaffolds such as that supported by Shank3B but this is entirely speculative.

### Strengths, Limitations and Conclusion

The current study had some significant strengths. The positive effects of the WBH protocol were observed in two different ASD-relevant mouse strains that are each characterised by different behavioural phenotypes at baseline. The findings are strengthened by the fact that fever range temperature for 4 hours was sufficient, in the absence of inflammation, to improve both positive and negative, strain-specific, behavioural phenotypes. The fact that improvements are not confounded by a generalised suppression of spontaneous activity, which occurs during the acute LPS-induced systemic inflammation, that would usually accompany the fever response, gives confidence that it is the elevated brain/body temperature *per se* that supports the improvements in behaviour. However, there are some significant limitations. Firstly, it was not possible to perform repeated testing of all behavioural parameters at all time points post-WBH and, indeed, it would also be beneficial to continue behavioural assessment well beyond 48 hours to assess whether there may be some long-term benefits to the treatment. More importantly, we provide only limited dissection of what cellular and molecular changes accompany or even facilitate behavioural improvements. We did observe dissociable patterns of immediate early gene expression (cFos) induced by LPS and WBH with a number of regions activated almost exclusively in WBH-treated animals. More detailed characterisation of these brain regions is required. Integrating the findings of (Reed *et al*., 2020) with previous humans studies (Curran *et al*., 2007; Grzadzinski *et al*., 2018), we propose that the presence or absence of improvements under hyperthermic conditions will likely depend on the initial etiology of the ASD, on the the behavioural symptoms examined and on the severity of inflammation and fever occurring. However given the potential of pyrogenic pro-inflammatory mediators such as IL-1β, IL-6 and prostaglandins to suppress motivation, mood, social activity, arousal and cognition (Saper *et al*., 2012), it remains plausible that the effects of elevated temperature could actually be more impressive if elevated temperature was achieved in the absence of inflammation. Despite the difficulties of such studies in human ASD cases, it is necessary to measure these parameters in order to come to more robust conclusions and to determine whether there is merit in pursuing the fever hypothesis for improvements in ASD.

## Funding and Acknowledgements

The work contained herein was supported by a grant from SFARI (Simons Foundation Award 422623 to CC). We thank Sharon Evans for initial discussions about whole body hyperthermia and Dr. Eugene Kiyatkin for helpful discussions on the design of thermocouple probes.

